# A microbiome meta-transcriptomics pipeline identifies a novel human neutrophil elastase inhibitor that protects the colonic epithelial barrier

**DOI:** 10.1101/2025.09.11.675725

**Authors:** Bojan Stojkovic, Matthijs Bekkers, Jake P. Violi, Brett A. Neilan, Simon Keely, W. Willem A. Donald, Carlos Riveros, Gerard E. Kaiko

## Abstract

Inflammatory Bowel Diseases (IBD) are lifelong conditions. Current therapeutic approaches target inflammatory signalling rather than improving barrier permeability or repair. The gut microbiome provides an exciting opportunity for novel drug discovery to leverage its role in healthy gut homeostasis. There is a clear need to identify bioactive molecules within the microbiota that could protect the intestinal barrier. Our group has developed a systematic pipeline using metatranscriptomic data to identify, produce, purify, and test microbial proteins in IBD, pinpointing multiple novel microbiota-derived proteins linked to disease activity. We identified a new microbiota protein (BMG-1), that specifically inhibits human neutrophil elastase, a pathogenic protease in IBD. This protease inhibition allows protection of the intestinal epithelial barrier from permeability and promotes epithelial healing. BMG-1 also reduces colon damage in a mouse model of colitis. These findings demonstrate the gut microbiota can specifically regulate the balance of protease/anti-protease activity in the colon, and this represents a novel therapeutic strategy for IBD.

## Introduction

Inflammatory bowel diseases (IBD) affect more than 7 million people worldwide. IBD, which includes ulcerative colitis (UC) and Crohn’s disease (CD), is a lifelong condition in which the intestinal barrier is recurrently damaged, leading to increased permeability of the gut, transepithelial migration of commensal microorganisms, excessive inflammation, and poor mucosal healing ^1–3^. The advent of biologic therapeutics has significantly improved the disease course, however, these drugs primarily aim to alleviate specific symptoms of IBD by dampening inflammation. This results in up to 50% of patients either failing to respond initially or losing response after 12 months, leading to a delay rather than prevention of the need for surgical intervention ^4,5^. There are multiple hallmark features in IBD that are predictors of disease course but are not targeted by current therapies. This includes impaired function of the epithelial barrier that leads to ‘leakiness’ or increased intestinal epithelial barrier permeability, as well as impaired mucosal healing. This leakiness allows for infiltration and/or exposure of immune cells to luminal contents and microbes, which is thought to perpetuate the chronic inflammatory response. This increased permeability is a predictor of both relapse and the initial onset of IBD ^6,7^. In addition, mucosal healing is considered one of the best predictors of long-term remission in IBD ^2,8,9^. The mass mucosal infiltration of neutrophils is another feature of IBD pathology and readily observed in histological biopsy specimens as part of the pathology diagnostic assessment ^10^. Uncontrolled neutrophil proteolytic activity leads to intestinal crypt damage and abscess formation ^1,11^. In addition, it has been demonstrated that the proteolytic enzymes that create this epithelial barrier damage are enriched in clinically active IBD patients ^12,13^, and known to precede the onset of IBD ^14^. Therefore, the development of therapies that protect the intestinal barrier by inhibiting these processes is an underexplored and innovative approach to IBD treatment.

The gut microbiome represents a vast and largely untapped resource of potential therapeutic molecules, and its dysbiosis has been consistently linked with IBD. The main signature of the gut microbiome change that occurs in both UC and CD is the overall loss of diversity and reduction in obligate anaerobic bacteria ^15,16^. Faecal microbiota transplantation (FMT), a method of transplanting healthy donor faeces, shows ∼30% endoscopic remission rate in UC patients with mild/moderate disease ^17–20^. However, exactly what microbes and what bioactive components within the FMT mixture are driving the positive response remains unclear. Furthermore, the stable engraftment of bacteria and the heterogeneous nature of this uncontrolled mixture remain a major problem with this approach. Far less research has gone into the bioactive proteins that healthy microbiota produce and how they may contribute to gut homeostasis or remission in IBD. Harnessing these molecular components could drive a much higher efficacy rate and would represent a more controllable and druggable pathway. Microbiome-derived molecules capable of protecting the epithelial barrier or promoting mucosal healing represent novel therapeutic avenues, targeting disease aspects currently not addressed by existing therapies.

In this study we developed a pipeline to identify microbial genes/proteins strongly linked to IBD diagnosis and disease activity, and then a platform to produce and purify these molecules. This pipeline used metatranscriptomic datasets as they enable identification of genes that are actively transcribed as opposed to metagenomics, which detects genes in dead bacteria, silent genes, and those with altered abundance but not expression. Therefore, changes in gene expression in metatranscriptomic data is more directly linked to changes in microbial protein activity. Our metatranscriptomic pipeline pinpointed uncharacterised novel microbial genes strongly associated with IBD disease activity indices and identified those likely to be secreted by bacteria at the intestinal barrier. These candidates were identified as potential microbiota drug targets in IBD. We then developed a platform to produce these microbial proteins and functionally test them for activity on human intestinal organoids. Using this pipeline we have identified a novel protein, which we named BMG-1, found to be a serine protease inhibitor. Our results demonstrate the ability of BMG-1 to specifically inhibit the human neutrophil elastase, which is elevated in IBD and responsible for intestinal mucosal damage ^21,22^. Importantly, we have also demonstrated a protective effect of BMG-1 on the epithelial barrier using colon organoids, and *in vivo* using the dextran sodium sulphate (DSS) model of colitis.

## Methods

### Bioinformatics pipeline

The publicly available datasets utilised in this study were sourced from the Human Microbiome Project 2 (HMP) substudy on IBD ^23,24^. This extensive longitudinal study encompassed data gathered from diverse cohorts, including 132 participants, comprising individuals with Crohn’s disease (CD) (n=67), ulcerative colitis (UC) (n=38), or non-IBD (n=27). Various analytical techniques, such as metagenomics (MTG) and metatranscriptomics (MTX), were conducted on collected stool samples. This study included bi-weekly sample collection over the one-year follow-up period. A total of 758 stool samples (371 CD, 210 UC, 177 non-IBD) from the dataset had paired MTG and MTX data.

A personal genomic reference scaffold was established for each patient by compiling all genomic references of the taxonomic units identified in the individual, using MetaPhlan2, from the individual’s MTG sequencing data ^25^. The identified taxa for each individual were collated and the corresponding reference genomes downloaded from National Center for Biotechnology Information (NCBI) to create a personalised genomic reference (PGR). The accuracy of this approach was confirmed by mapping all MTG reads to the scaffolds on a per sample basis. Following read quality assessment using FastQC, we applied Cutadapt for adapter trimming. MTX reads were then mapped to the genomic reference using Bowtie2 ^26^. Individual (time point) sequenced data with less than 0.5 million read pairs (MTG) or single end reads (MTX) were excluded. As a quality control of the completeness of the PGRs, unmapped reads from all individuals with more than 10% of unmapped reads were mapped to the global genomic reference (GGR), a reference constructed from all species from all subjects used in the study. Low numbers of unmapped reads on the GGR from PGRs with a moderate or high percentage of unmapped reads indicated incompleteness of the PGR and prompted us to manually expand these specific PGRs with genomes of missing species and strains. For each individual, a ratio was calculated by taking the fraction of unmapped reads that mapped to the GGR and dividing this by the fraction of initially unmapped reads and plotted per individual. For individuals with incomplete personalised genomic references, the cut-off ratio of 0.2 was used for updating the personal reference. All sequencing data from individuals with expanded PGRs were mapped again to the updated PGRs. After this step, all individuals had less than 10% of unmapped reads in MTX, and less than 15% in MTG.

To consistently identify putative expressed proteins for all individuals for the quantitation of mapped sequence data, all open reading frames (ORFs) were identified *de novo* from the GGR using Prokka, a tool designed for finding ORFs in prokaryotes ^27^. Features such as rRNA, tRNA and tmRNA, that do not encode for protein sequences, were identified using ViennaRNA and were not relevant to our pipeline, excluding them from the annotation of our GGR ^28^. Then, gene clustering was performed based on amino acid sequence similarity, grouping ORFs into two categories: 100% and 98% amino acid identity to enable the investigation of both unique predicted microbial proteins and those sharing a high degree of similarity but may only differ by 2% of amino acids perhaps among similar strains of the same species ^29^.

Mapped sequence data was then counted towards these genomic annotations using FeatureCounts ^30^. A minimum cut-off value of 5 reads per gene was applied to ensure robustness. Multiple mapping reads, of which we kept the top 5 mapping locations, were counted using fractional counts to all their mapping locations. This approach expanded the number of genes with counts for differential expression analysis without disproportionately overrepresenting genes with very similar sequences across multiple species.

For disease diagnosis, two separate contrasts were used in the DE analysis between disease groups: CD vs non-IBD and UC vs non-IBD. To assess differences in microbial gene expression concerning disease activity, the IBD (CD and UC) samples were stratified based on faecal calprotectin (FC) levels, the most clinically common molecular biomarker for disease activity. For this stratification, samples from UC and CD patients were aggregated into two groups: low disease activity and high disease activity, as defined by an FC threshold of 200 μg/g, which is within the recommended range for patient classification ^31^. Overall,148 samples contained FC data, of which 96 samples were classified as low disease activity, and 52 samples as high.

The statistical models used in the DE analysis, implemented in the R packages DESeq and VariancePartition, have distinct statistical assumptions about the underlying data distributions, leading to different analytical approaches to increase the confidence in any resulting DE gene hits. Their unique assumptions result in model-specific advantages in handling zero count reads and normalisation ^32–34^. To be conservative, we applied both statistical models to our DE analysis, ranking the genes based on p-value and fold change. For each contrast (CD vs non-IBD, UC vs non-IBD, and low vs high activity), 5000 top-ranked genes were compared, and overlapping common genes were analysed further. Following the DE analysis, we aimed to prioritise genes that are more likely to be biologically relevant. We hypothesised that gene candidates would be of greater biological interest if they were detected in a substantial portion of the sample population, and we set a presence threshold of 30%. This indicated that the gene had to be detected as being expressed in at least 30% of the samples in at least one of the groups in the contrast. The basis of the model and code is included in https://github.com/BioAI-CR/BIO18025-FunMG.git

### Gene candidate selection

Differentially expressed gene lists were further filtered by identifying predicted secreted proteins as we hypothesised that secreted proteins or extracellular microbial proteins that contained signal peptides would be more likely to exert functional interaction with the host. This filtering step was based on multiple prediction algorithms that identified signal peptides on amino acid sequences EffectiveDB and SignalP ^35,36^.

Additionally, our selection process for practical laboratory production involved the assessment of gene sizes, considering limitations for the heterologous expression and purification of large proteins in *E. coli*. Genes larger than 1500 base pairs (∼50 kD) were excluded. Next, gene sequences were checked for the presence of restriction enzyme sites that could interfere with cloning. After these filtering steps following DE analysis, a total of 14 genes were chosen. We aimed to purify a panel of ten IBD-linked microbial proteins for screening to identify those that have functional effects on epithelial barrier permeability and repair.

### Recombinant protein synthesis

The design of the DNA constructs, *in silico* cloning and multiple sequence alignments were performed using SnapGene (GSL Biotech LLC). Gene sequences were codon optimised for *E. coli* and modified by the addition of *SapI* restriction site to facilitate ligation into an expression vector. The final gene fragments were commercially synthesised (Integrated DNA Technologies), *SapI* digested and ligated using Electra system according to instructions (Atum). Two rhamnose inducible expression vectors were utilised: pD861-His or pD861-HisMBP, with N-terminal 6xHis or 6xHisMBP fusion tag, respectively. The correct gene insertion was confirmed using colony PCR and Sanger sequencing (Australian Genome Research Facility). The positive clones were grown to OD600=0.4, induced using 4mM Rhamnose, and cultured overnight at 23°C for optimal protein expression. The bacterial pellets were freeze-thawed, lysed using the combination of B-PER™ reagent treatment (Thermo Fisher Scientific) and sonication (30% power and pulse for 2min). The insoluble fraction and debris were removed by centrifugation and the supernatant containing the recombinant protein was applied to Immobilized Metal Affinity Chromatography (IMAC) columns and purified according to manufacturer’s instructions (Cytiva). The HisMBP tag was cleaved with His-tagged HRV-3C Protease (Sigma) at the specific cleavage site present on the vector and removed along with the protease with a secondary IMAC step. After purification, samples were further processed using Pierce High-Capacity Endotoxin Removal Spin Columns, following manufacturer’s instructions (Thermo Fisher Scientific). Protein concentration and purity were assessed using Pierce™ BCA Protein Assay Kit (Thermo Fisher Scientific) and 4–15% Mini-PROTEAN™ TGX Stain-Free SDS-PAGE (BioRad) and Pierce™ Chromogenic Endotoxin Quant Kit (Thermo Fisher Scientific). The presence of His tagged BMG-1 (HisBMG-1) was confirmed by Western Blot using the anti 6xHis Tag Polyclonal Antibody (Thermo Fisher Scientific).

### Mass spectrometry

For characterisation of the microbiota-derived protein, HisBMG-1, we performed native mass spectrometry. Recombinant HisBMG-1 was buffer exchanged into 100 mM ammonium acetate (pH ∼7.5) using Zeba spin desalting columns (Thermo Fisher Scientific). Protein concentration was adjusted to >1.5 µg/mL for all experiments. To assess interactions with human HNE, HisBMG-1 was incubated with HNE at molar ratios ranging from 1:2.5 to 1:10 (HNE:HisBMG-1) for up to 1h at room temperature prior to MS analysis. Mass spectra were acquired using Q Exactive UHMR Hybrid Quadrupole Orbitrap (Thermo Fisher Scientific), under previously described conditions optimised for native MS ^37^. Spray voltage was set to 0.8 -1.2 kV with the heated capillary entrance being set to 200 °C. All analyses were performed in positive ion mode using a mass range of 2000 – 6000 *m*/*z*. Samples were introduced by direct infusion using static nanospray emitters. Deconvoluted mass spectra were obtained using UniDec, and charge state assignments were manually confirmed based on isotopic spacing and m/z shifts between adjacent charge states. Theoretical masses were calculated from protein sequences and compared to deconvoluted peaks.

### Human neutrophil enzyme isolation

Blood was collected from healthy volunteers as approved by the Hunter New England Health human research ethics committee (2020/ETH03303). Written informed consent was obtained from the study subjects. The whole blood was diluted 2:1 with 3% dextran saline solution to remove the red blood cells. The top layer was diluted with an equal volume of 2% fetal calf serum (Thermo Fisher Scientific) in PBS. Next, 8 ml mixture was layered on 4 ml Lymphoprep^TM^ media, and processed according to instructions (StemCell Technologies). The cell pellet was treated with the red blood cell lysis buffer (150 mM NH4Cl, 10 mM NaHCO3, 0.1 mM EDTA) for 2-5 min. After washing with Hank’s Balanced Salt Solution (Thermo Fisher Scientific), cells were resuspended in DMEM media (Thermo Fisher Scientific), counted, and purity of neutrophils was assessed using May-Grunwald Giemsa Staining and cytospin (>90%). Neutrophil cell lysate was prepared by one freeze thaw cycle, sonication (30% power and pulse for 2 min) and final bead beating step (30 Hz, 5 min). Debris was removed by centrifugation and cell lysate was aliquoted and stored at -80°C.

### Culturing of human colonic organoids

Human colonic organoids were provided by the Digestive Health Biobank as approved by the Hunter New England Health human research ethics committee (2020/ETH03303). Written informed consent was obtained from the study subjects. Organoid lines were isolated from colonic biopsies from non-IBD patients, expanded and biobanked ^38^. Organoids were cultured in 24 well plates, in three-dimensional 40 µl dome structures in Matrigel (BD Biosciences) and expanded using our well-established L-WRN stem cell conditioned media (CM) system ^39–42^. For the expansion of organoids CM was further supplemented with SB 431542 ALK inhibitor (TGFβi, 10µM, Abcam), Y-27632 dihydrochloride Rho kinase inhibitor (ROCKi, 10 µM, Abcam), SB 203580 p38 MAPK inhibitor (p38i, 5 µM, Abcam) and Primocin (1mg/ml, InvivoGen) ^38^.

### Intestinal epithelial barrier permeability assay

After organoid culturing, *in vitro* epithelial barrier permeability assays were conducted by disaggregating the 40 µl organoid domes into single cells using Trypsin-EDTA (Thermo Fisher Scientific) for the formation of differentiated cell layers producing mucous ^42^. After enzymatic digestion, cells were plated on a 96 well transwell system, containing a 0.4 µm pore polyester membrane (Stem Cell Technologies), in a 1:5 ratio. The cells were incubated with the complete CM, on both basal (120 µl) and apical side (40 µl). Upon reaching confluency, CM was removed from the apical side, to create an Air Liquid Interphase (ALI) culture that was differentiated for 2 or 10 days, depending on the experiment. On the final day, the cells were washed with DMEM media (Thermo Fisher Scientific) to remove serum and subsequently treated on the basolateral side for 6 h with the experiment specific molecules. These included neutrophil lysate (1:2 dilution in DMEM), or recombinant human neutrophil elastase (HNE, 500nM, Abcam), in combination with or without microbial proteins of interest. Elafin (R&D systems) was used as a positive control. Fluorescein Isothiocyanate (FITC) conjugated Dextran 4,000 MW was then added to the apical side and epithelial permeability was measured in the basolateral side over a 6 h time-course by assessing FITC movement across the layer as an established surrogate of permeability. The fluorescent signal was measured using CLARIOstar microplate reader, and FITC Dextran concentration was calculated using 4 parameter logistic standard curve.

### Wound repair and cell death assay

Our *in vitro* wound repair assay was conducted on primary human colonic organoid cells. Organoids were passaged as described above and seeded into monolayers in sterile 96 well flat bottom tissue culture plates at 1:4 ratio and grown to 90% confluency. Cells were then incubated for 24 h before they were mechanically scratched using the Essen Bioscience IncuCyte® WoundMaker - 96-pin wound making tool (Essen Bioscience Inc.). The cells were then washed three times with DMEM media (Thermo Fisher Scientific) to remove traces of serum and cell debris; and treated with 100 nM HNE premixed with microbiota proteins or controls at various concentrations in differentiation media (advanced DMEM/F-12 (Thermo Fisher Scientific), supplemented with 2 mM L-glutamine, 0.5mg/ml penicillin/streptomycin, 50ng/ml hEGF and 10uM Y-27632). Wound healing was monitored at 6 h intervals and analysed over 72 h using the automated IncuCyte® cell migration image capture and analysis software (Essen BioScience, Inc.). For the purpose of cell death assay, cells were cultured and seeded as described above. Upon reaching 100% confluency, cells were washed twice and treated with 250 nM HNE premixed with microbiota proteins or controls at various concentrations, and cultured in differentiation media for 24 h. Images were taken every 3 h and confluency was measured using the automated IncuCyte® cell confluency image capture and analysis software (Essen BioScience, Inc.). At the 24 h time-point, cell metabolic activity was measured using CellTiter 96® Non-Radioactive Cell Proliferation Assay (MTT), according to manufacturer’s instructions (Promega).

### Cell free protease assay

The protocol was adopted from Mkaouar *et al*. ^43^. Briefly, proteases (neutrophil lysate, purified enzymes) and microbiota proteins of interest were mixed in a flat bottom 96 well plate after which the MCA substrate (RnD biosystems) Mca-RPKPVE-Nval-WRK(Dnp)-NH2 was prepared in assay buffer (20 mM Tris-HCl, 50 mM NaCl Buffer, pH 8) at a final concentration of 10 µM. MCA substrate was added and fluorescence was measured (excitation 320-15nm, emission at 405-20nm) at different timepoints at 37℃. Raw relative fluorescence unit (RFU) values were normalised to the background control, and expressed as percentage enzyme activity.

### DSS murine colitis model

C57BL/6 male mice (6 to 8 weeks old) were obtained from Australian BioResources. All mice were kept at room temperature with 12-h light/dark cycles and free access to food and water. All procedures were approved by the Animal Care and Ethics committee at the University of Newcastle. After the one-week acclimatisation period, mice were treated with 2% DSS dissolved in drinking water for 6 days, to induce colonic inflammation. Weights were monitored daily. HisBMG-1 (10mg/kg) or vehicle control were administered intrarectally every 2 days. DSS treated mice groups were euthanized on day 8, due to standard ethical requirements at 20% weight loss. The vehicle treated mice groups endpoint was day 10. Faecal samples were collected at endpoint for protease activity measurement. Samples were weighed and dissolved in 1 ml resuspension buffer (50 mM Tris, 150 mM NaCl, pH 7.5) per 100 mg, homogenised by bead beating, centrifuged at 16,000 x *g* for 10 min to remove the debris. Total protein content was quantified in the supernatant, using BCA assay and samples were diluted to 200 µg/ml. Two microliters per sample was then mixed with a broad-spectrum protease substrate (RnD biosystems) Mca-RPKPVE-Nval-WRK(Dnp)-NH2, and protease activity assay was performed as described above. Colon length was measured, and colon tissue was harvested for histological assessment of damage. The DSS induced epithelial damage was evaluated based on colon histopathology by H&E and PAS. The percentage of normal colon tissue that had no histological mucosal ulceration, no goblet cell loss, or loss of crypt architecture was measured as previously described ^44^.

### Statistical analysis

GraphPad Prism (GraphPad Software) was used for all statistical analysis. The data from the epithelial permeability assay, wound repair/cell confluency assay and cell free protease assay was analysed using repeated measures two-way ANOVA or Friedman test and Dunnett’s multiple comparison test. *In vivo* DSS mouse model data was analysed using ordinary one-way ANOVA and Dunnett’s multiple comparison test.

## Results

### Systematic analysis of the gut microbiome transcriptome identifies microbial genes associated with UC and CD diagnosis as well as disease activity

Microbial gene expression is often overlooked in the gut meta-omic analysis of IBD, as it has mainly prioritised taxonomic profiling of species or identifying changes in broad categories of genes measured by abundance through metagenomics. Using our bioinformatic pipeline, and the largest publicly available metatranscriptomic datasets from the Human Microbiome Project 2 (HMP) substudy on IBD ^23,24^, we aimed to identify specific microbial gene/protein candidates linked to IBD. We then prioritised a subset of the genes for further functional testing in the laboratory (Figure 1A). We utilised an *E. coli*-based system for gene candidate cloning and protein overexpression. The protein products of top gene candidates were purified and tested for their functional properties using *in vitro* and *in vivo* models of intestinal disease (Figure 1B).

**Figure 1.**
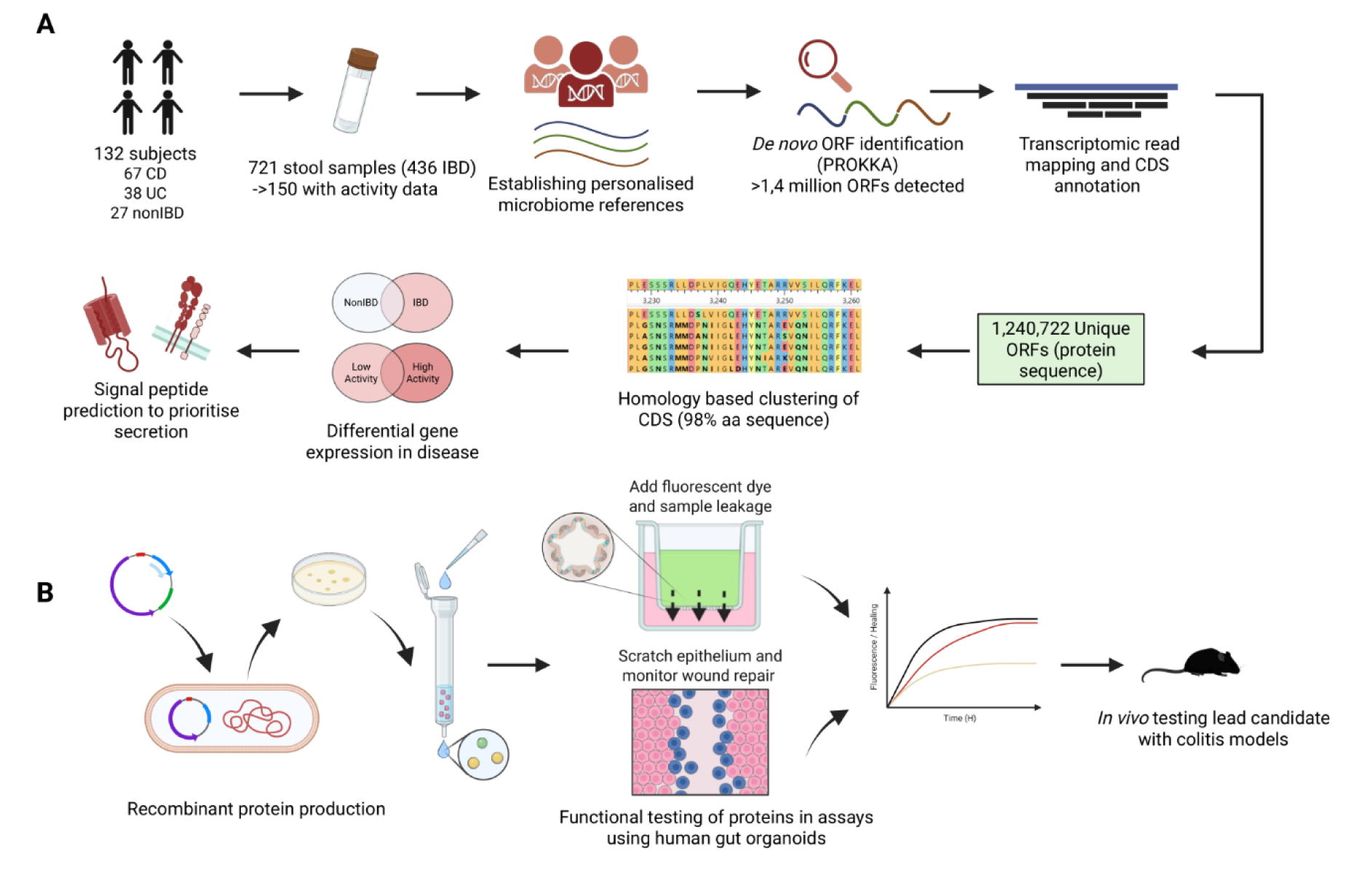
Overview of pipeline for the identification, synthesis and testing of novel microbial genes associated with IBD disease status and disease activity. (A) Overview of bioinformatic pipeline for identification of IBD linked genes (expanded in Figure S1). Personal genomic references were established for each subject from publicly available metagenomic sequencing data (HMP) using MetaPhlan2. Identification of open reading frames (ORFs) was conducted on the genomic reference followed by mapping of metatranscriptomic (MTX) reads. Genes were then clustered based on sequence homology. A differential expression (DE) analysis was performed on the following comparisons, CD vs non-IBD, UC vs non-IBD, and low vs high disease activity. Further gene candidate filtering was performed, based on the secretion prediction, statistical significance, log fold change and downstream wet lab compatibility. Finally, top candidate genes were selected for further analysis. (B) Overview of the gene candidate testing pipeline. Gene fragments were cloned into an *E. coli* vector expression system. Proteins were synthesized and purified for functional testing using *in vitro* human intestinal organoid-based assays and *in vivo* murine colitis model.

The summary of the bioinformatic gene filtering is shown in Figure 2A. The transcriptomic sequencing reads were mapped with an efficiency of ∼80-90%, to the personalised genome references, which was further improved upon additional mapping to the global reference (as described in methods; Figure S1-A). The *de novo* ORF identification was then conducted, producing 1.43 million ORFs across all metatranscriptomic samples. Genes were further filtered for translational redundancy by gene clustering, resulting in 1.1 million ORFs (Figure 2A). The gene clustering was based on the amino acid homology of >98% and protein length similarity >98% (Figure S1-B), aiming to identify the groups of highly homologous genes likely among the strains of the same species with similar functions. To identify microbial genes differentially expressed in disease, the differential gene expression (DE) analysis was carried out for three pairwise comparisons independently: UC vs nonIBD, CD vs nonIBD and low vs high disease activity. The disease activity classification was determined using the common clinical faecal calprotectin threshold of 200 µg/g, which stratified the samples into two distinct categories (Figure S1-C). Genes were filtered using the p-value, and a log2-fold difference of 2 (4-fold) (Figure 2B). In addition, a disease relevance filter was applied which only included genes detected in at least 30% of patients from either classification group. This threshold was chosen based on the observation that the total number of detected reads per sample stabilises when approximately 30% of samples are considered (Figure S1D-E). Due to the nature of microbiome data two statistical models for DE were used, DESeq2 and Variance Partition. We only considered DE genes detected by both statistical models. The DE analysis for all three comparisons (UC vs nonIBD, CD vs nonIBD and low vs high disease activity) was performed on both 98% and 100% gene clusters, producing a total of 24,680 differentially expressed microbial genes (Figure 2A). We then carried out the shared feature analysis to identify genes present across both statistical methods and clustering groups, resulting in 797, 182, and 1431 genes in low vs high (Figure 2C), CD vs nonIBD (Figure S1-F) and UC vs nonIBD comparison (Figure S1-G), respectively. Importantly, functional annotation of differentially expressed genes showed that more than half of the genes are of unknown function, when using both clustering of orthologous genes (Figure 2D-F), or categorisation based on the enzyme commission numbers (Figure S1H-J).

**Figure 2.**
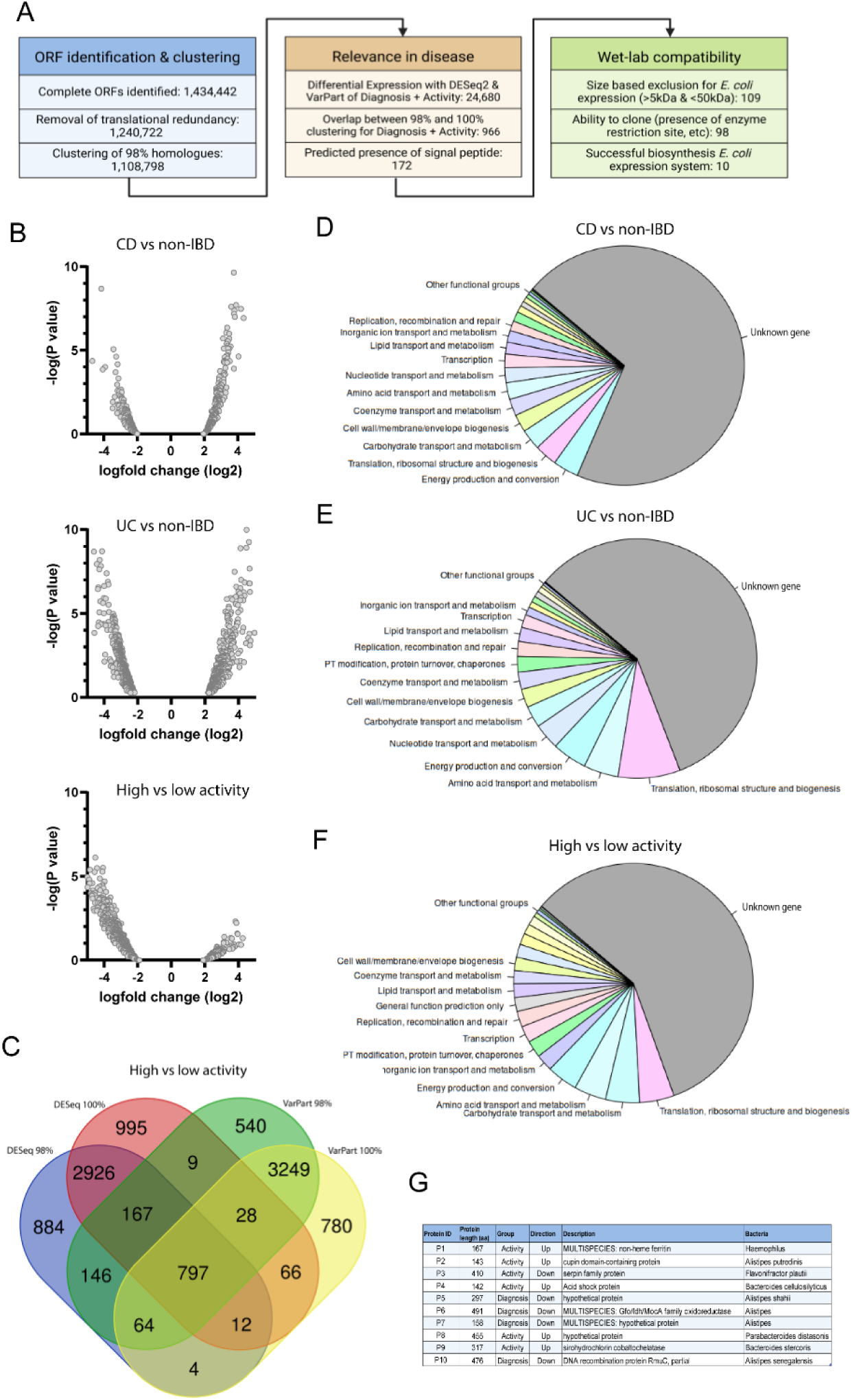
Characterisation and prioritisation of differentially expressed genes in the metatranscriptomic analysis of the gut microbiome in IBD. (A) Discovery pipeline for top candidate microbial gene selection in IBD. Genes were filtered and prioritised based on several features such as homology clustering, magnitude of differential expression, likelihood of secretion and wet lab compatibility for cloning. (B) Volcano plots of top DE genes generated by 98% clustering analysis for 3 comparisons CD vs non-IBD (752 genes), UC vs non-IBD (2134 genes), and high vs low disease activity (1177 genes). (C) Venn Diagram on gene lists from 4 comparisons; high vs low disease groups for the Variance Partition (VarPart) and DESeq2 model and two clustering strategies (98 and 100%) to find DE genes common to all. (D, E, F) Pie charts representing Clusters of Orthologues Genes (COGs) from top DE genes. (G) 10 microbial genes selected for protein synthesis and functional testing.

Subsequent filtering by prioritising genes with predicted signal peptides, yielded 172 DE gene candidates predicted to be secreted. We then employed additional gene selection to accommodate practical considerations of *E. coli* protein production. Specifically, genes encoding proteins below 5kD and over 50kD, and genes with a specific restriction enzyme binding site, were excluded, resulting in a final list of 98 genes (Figure 2A), of which 10 were successfully synthesised and sufficiently purified in the laboratory and selected for further testing (Figure 2G).

To examine the taxonomic profile among the top differentially expressed genes from metatranscriptomic analysis (LFC > 2 and P-value < 0.0001, ∼9000 genes), we listed the top 120 species for all disease comparisons in both 98% and 100% clustering strategies (Figure S2). This selection covered a high number of species that have previously been associated with IBD, including consistently reported taxa such as *Faecalibacterium prausnitzii, Roseburia hominis, Roseburia inulinivorans and Veillonella parvula* ^45–48^. In addition to exploring taxonomic features in our analyses, we investigated the potential of our DE genes as predictive biomarkers. We performed Random Forest classifier-based feature selection and pairwise feature combinations. ROC curves and area under the curve (AUC) scores for each contrast group are shown in Figure S3. The Random Forest classifier provided the ROC curve performance of a multi-feature classifier from which a Mean Decrease Accuracy ranking was generated. For the pairwise analysis, CD vs non-IBD, UC vs non-IBD and high vs low activity showed AUC values of 0.83, 0.9, and 0.85, respectively (Figure S3A-C). For the Random Forest classifier, AUC values of 0.96, 0.99, and 0.83 were obtained for CD vs non-IBD, UC vs non-IBD and high vs low comparisons, respectively (Figure S3D-F). This data strongly suggests a high accuracy for the expression level of bacterial protein-coding genes, especially in pairs, to predict both UC and CD diagnosis, as well as IBD disease activity.

### Functional screening of top gene candidates identified BMG-1 as a microbial protein with protective properties on colonic epithelial cells

Identifying microbial proteins with the ability to protect the colon epithelium from damage or mediate epithelial repair is of great importance, as mucosal healing is critical to long-term remission in IBD ^2,8,9^. Therefore, we employed a human gut organoid epithelial barrier assay to test the ability of microbial proteins to either induce mucosal damage, or protect the epithelial surface. As neutrophils are the most abundant immune cells elevated in IBD patient mucosal biopsies, and a known damaging agent for the intestinal barrier, we used neutrophil contents (lysate) to induce damage in order to assess the ability of microbial proteins to protect against damage (Figure 3A). Among the 10 tested microbial protein candidates, microbial protein P3 (referred to as BMG-1) exhibited the greatest, dose-dependent improvement in epithelial barrier permeability (Figure 3B). We analysed BMG-1 gene expression in the metatranscriptomic data and found it to be highly upregulated (∼20 fold) in IBD faecal samples with low disease activity and reduced in states of high disease activity (Figure 3C). This observation remained consistent even after excluding samples without observed gene expression for BMG-1 (Figure 3D). The BMG-1 gene was expressed in a high portion of samples with reads detected in 70%, 79%, and 56% of samples from nonIBD, low, and high disease activity group, respectively (Figure 3E). Next, we investigated BMG-1 gene expression in IBD samples and its correlation to the disease activity biomarker faecal calprotectin and as expected found a strong inverse correlation (Spearman r=- 0.32, P= 0.001) (Figure 3F). Even after including nonIBD samples, this correlation decreased but remained significant (Spearman r=-0.-16, P= 0.0242) suggesting a role of this protein in disease activity in IBD (Figure 3G). Lastly, we correlated BMG-1 gene expression with relative abundance of *Flavinofractor plautii*, one of its predicted bacterial species of origin, within the HMP2 cohort and found a strong correlation between the gene and taxa (Spearman r= 0.77, P= 0.0001) (Figure 3H).

**Figure 3.**
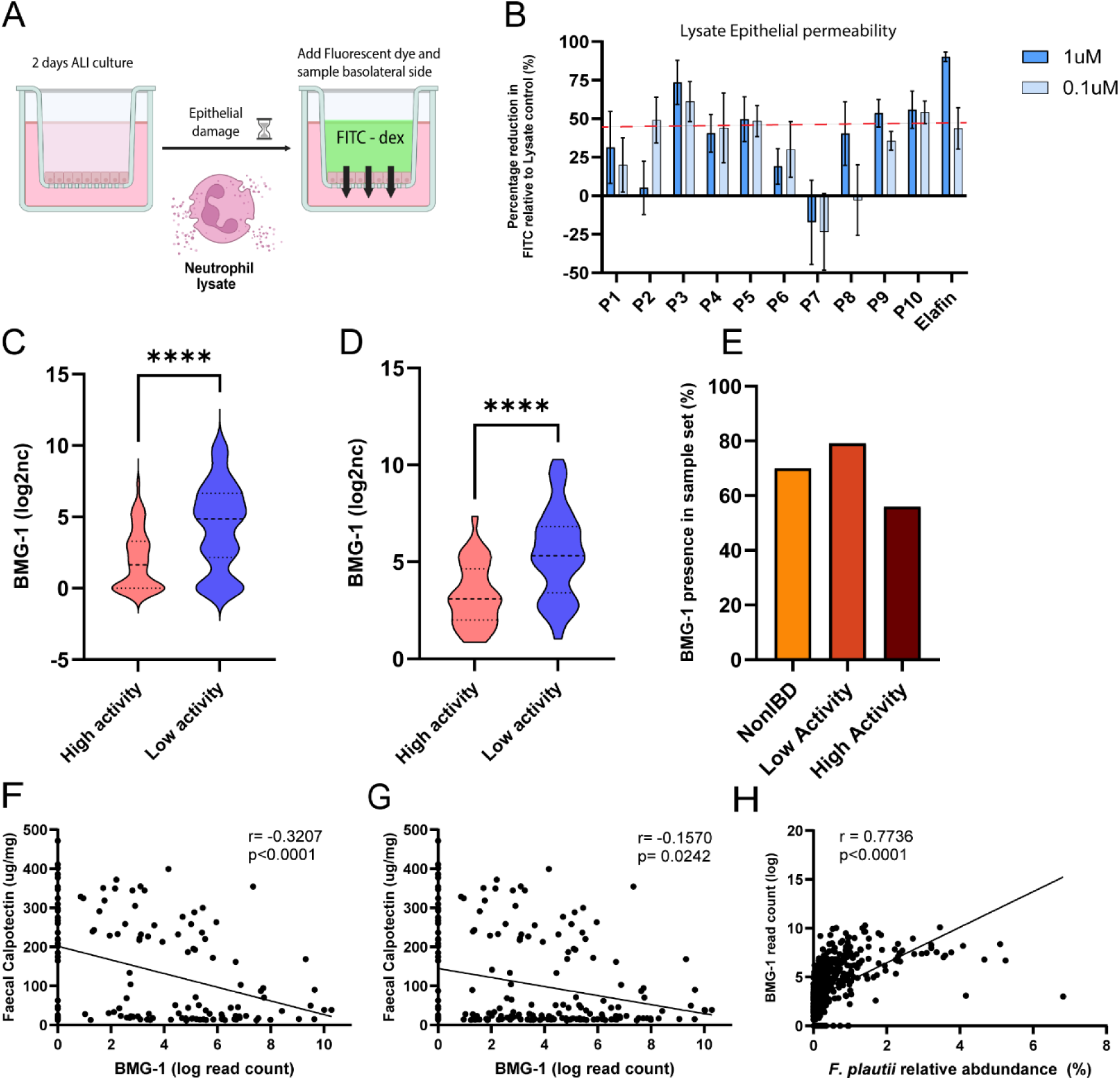
Functional screening of top 10 microbial gene candidates and BMG-1 identification. (A, B) Functional testing of synthesised microbial proteins from top prioritised genes in an *in vitro* human organoid assay. Effects of 10 proteins on epithelial permeability was assessed on air liquid interface colon organoids pretreated with neutrophil lysate (NeutLys). (C, D) Violin plots show differential expression analysis results (log fold change) of BMG-1 RNA expression between the high and low IBD disease activity groups. (C) Graph represents all samples from disease activity analysis, or (D) samples that display zero BMG-1 expression or below detection were excluded. Groups were compared using parametric t test p<0.001 (****). (E) Percentage samples with detected BMG-1 gene expression. (F, G, H) Correlation of BMG-1 gene expression in HMP cohort samples with faecal calprotectin (FC) in IBD (F) (moderate correlation, Spearman r=-0.320, P<0.0001), FC of IBD + nonIBD (G) (low correlation; Spearman r=-0.157, P= 0.0242), and relative abundance of *F. plautii* (H) (strong correlation; Spearman r= 0.77, P<0.0001).

### BMG-1 is a potent neutrophil protease inhibitor

BMG-1, identified via functional screening as a molecule with colonic barrier protective properties, was then further validated for its ability to protect the colon epithelial layer, over a 6 h time-course (Figure 4A). BMG-1 was produced and purified with and without a His-tag, HisBMG-1 (N-terminal 6xHis-tag) or BMG-1, and both were tested throughout our experiments. A clear dose effect was observed, with the highest dose of HisBMG-1 significantly reducing the epithelial permeability over this time course experiment (Figure 4A). This effect was replicated using BMG-1, without the 6xHis-tag (Figure S4-A), and was comparable to elafin, a known mucosal protease inhibitor used as the positive control (Figure S4-B). As BMG-1 prevented damage to colonic epithelial barrier cells that was induced by neutrophil contents, we hypothesised that BMG-1 may act through inhibiting neutrophil proteases, which are the main contents of neutrophil granules. To test this, we assessed the HisBMG-1 inhibitory function in a cell-free protease assay, using human neutrophil lysate and a fluorescent protease substrate. A statistically significant dose dependent inhibition of neutrophil protease activity was observed at 40min, 1h, 2h and 3h (Figure 4B). We then went back to our metatranscriptomic datasets and explored the homology clustering results around BMG-1 amino acid sequence and identified BMG-2, a highly similar protein characterised by 13 amino acid substitutions compared to BMG-1, with an overall similarity of 98% (Figure S4-C). We successfully cloned and purified this microbial protein and tested its ability to likewise inhibit protease activity. Interestingly, HisBMG-2 completely failed to suppress neutrophil protease activity (Figure 4C), highlighting the sequence specificity of BMG-1 and strongly supporting our decision to analyse genes from the 100% and 98% amino acid clustering homology lists separately in our bioinformatics pipeline. The HisBMG-1 neutrophil protease activity was successfully replicated using untagged BMG-1 (Figure S4-D) and elafin also achieved this inhibition (Figure S4-E).

**Figure 4.**
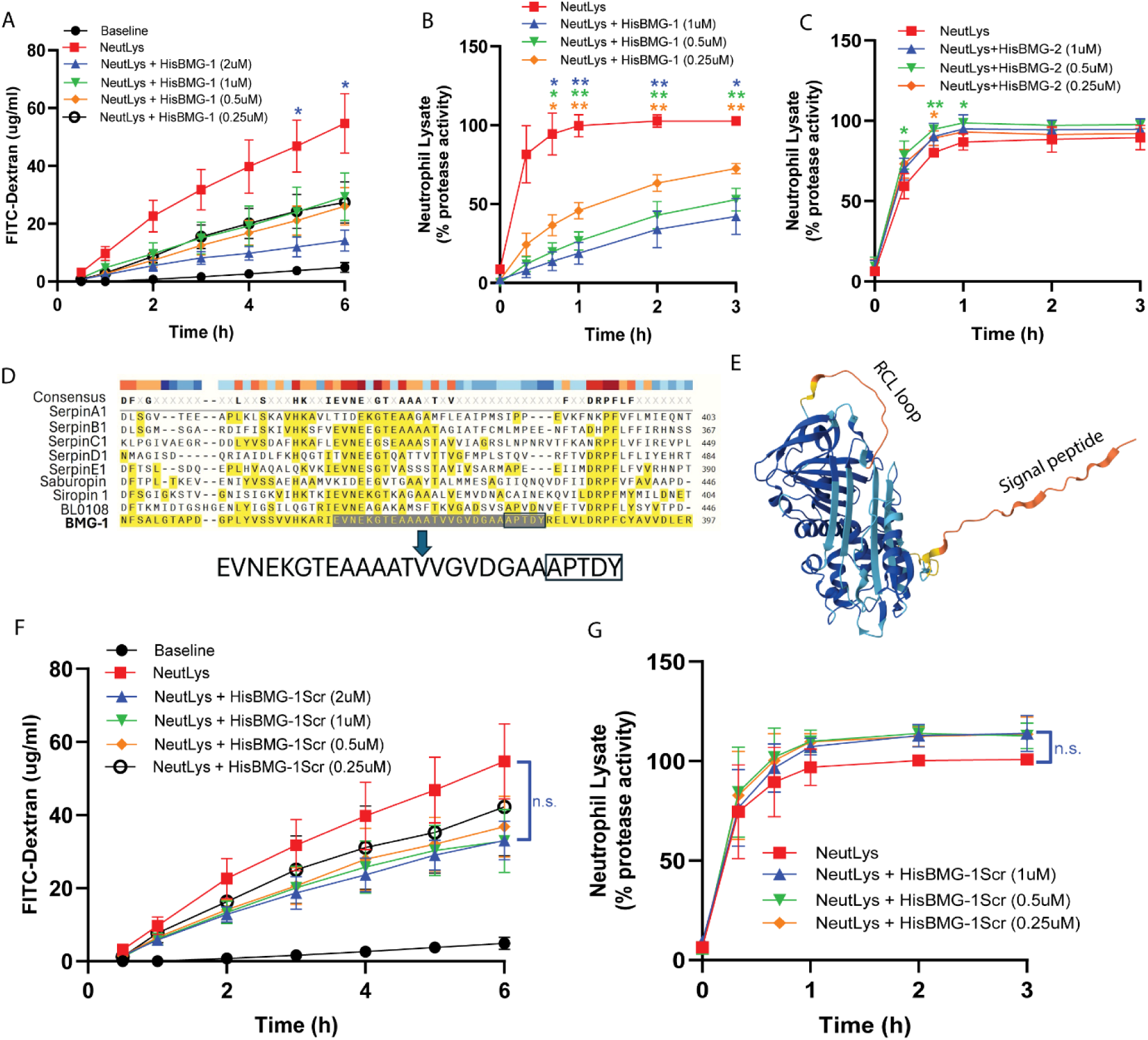
BMG-1 inhibits human neutrophil protease activity and protects the colon epithelium from neutrophil-induced damage. (A) Permeability of air-liquid-interface colon epithelium pretreated with neutrophil lysate (NeutLys) ± HisBMG-1, measured over 6h time-course. (B-C) The effect of HisBMG-1 (B) and HisBMG-2 (C) on neutrophil lysate protease activity in a cell-free protease assay. (D) Protein sequence alignment between BMG-1 and human serpin A1 - E1, *Eubacterium saburreum* Saburopin, *Eubacterium siraeum* Siropin and *Bifidobacterium longum* BL0108, highlighting the reactive centre loop (RCL) sequence and the scrambled region APTDY. (E) AlphaFold predicted 3D structure of unprocessed BMG-1 showing the RCL loop and the signal peptide extending from the main body of the protein. (F) Permeability of colon epithelium pretreated with neutrophil lysate ± HisBMG-1Scr, measured over 6h time-course. (G) The effect of HisBMG-1Scr on the neutrophil lysate protease activity in a cell-free protease assay. All data is presented as mean ± SEM, n=6 (A), n=4 (B-C), n=3 (F-G). p-adj<0.05 (*), p-adj<0.01 (**), Treatments were compared using repeated measures two-way ANOVA and Dunnett’s multiple comparison test. Significance between NeutLys and HisBMG-1//HisBMG-2/HisBMG-1Scr is marked by an asterisk matching the dose colour.

In order to better understand the functional properties of BMG-1, we performed *in silico* multiple alignments of the BMG-1 protein sequence, with 5 known human genes that function as serine protease inhibitors, belonging to the serpin superfamily: Serpin A1, Serpin B1, Serpin C1, Serpin D1 and Serpin E1; and three gut bacterial serpin protease inhibitors from *Eubacterium saburreum* Saburopin ^49^, *Eubacterium siraeum* Siropin 1 ^43^ and *Bifidobacterium longum* BL0108 ^50^ (Figure 4D). While overall homology was very low (below 30%), we identified a sequence of 26 amino acids in a domain in the C terminal region of the protein with significant homology across most of these molecules (Figure 4D). This suggested a relatively similar domain among these protease inhibitors, which may be the key enzyme active domain. Further 3D modelling of BMG-1 using AlphaFold 3 ^51^, revealed a typical serpin-like structure with 9 α-helices, and a central β-sheet forming the main body of the protein. Albeit with relatively weak confidence, the reactive central loop (RCL) region and the signal peptide were predicted in this structure and are observed protruding from the main body of the protein (Figure 4E). Importantly, we found that this RCL region lies within the conserved 26 amino acid C terminal domain, suggesting that this region may be critical for the protease inhibition function. To test these *in silico* predictions more directly, we then engineered four mutated versions of HisBMG-1, by scrambling the domains of the RCL region, and successfully purified 4 versions of a HisBMG-1 scrambled control protein (Figure S4-F). Mutations in the RCL region abrogated the protease inhibition in all four proteins to varying extents, with scrambled-1, -2, and -4 having the greatest reduction in activity (Figure S4-G). Protein Scr2 was selected for all downstream experiments and will be referred to as ‘HisBMG-1Scr’ throughout the text. In contrast to HisBMG-1, HisBMG-1Scr failed to protect the colon epithelial cells from damage by neutrophil granules (Figure 4F), and also lost the ability to inhibit the neutrophil proteases in the cell-free protease assay (Figure 4G). Together, functional and structural analysis demonstrates that BMG-1 is a potent neutrophil protease inhibitor, which is also reflected in its ability to block neutrophil-induced damage on the colon epithelium *in vitro*. In addition, we identified the critical region for BMG-1 function, pinpointing the RCL domain at the C terminus of the protein.

### BMG-1 specifically inhibits the human neutrophil elastase

We next aimed to identify the specific enzymes in the neutrophil protease contents that are targeted by BMG-1. We tested the 3 main enzymes known to be present in neutrophil granules, Cathepsin G, Proteinase 3, and human neutrophil elastase (HNE) using the cell-free protease assay. The activity of Cathepsin G and Proteinase 3 were unaffected by BMG-1 (or HisBMG-1). BMG-1 strongly and specifically inhibited HNE, a protease that is known to be linked to IBD pathogenesis (Figure 5A and Figure S5A-C).This effect was completely abrogated in HisBMG-2 (Figure 5B) and the scrambled control HisBMG-1Scr (Figure 5C), suggesting the importance of sequence specificity and the RCL loop in mediating this inhibition of HNE.

**Figure 5.**
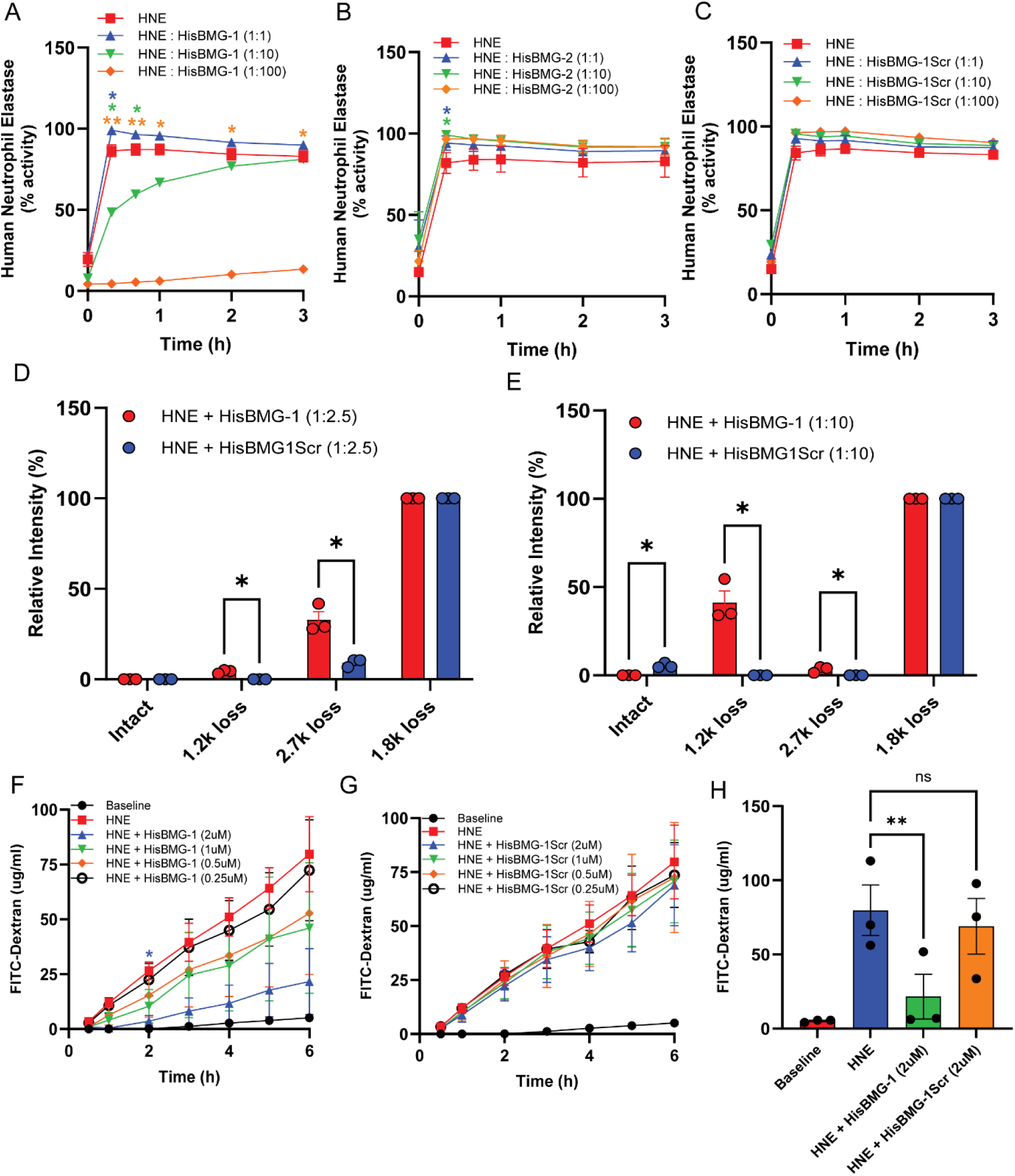
BMG-1 inhibits human neutrophil elastase (HNE) and protects the colon epithelium from elastase-induced damage. (A-C) The effect of HisBMG-1 (A), HisBMG-2 (B) and HisBMG-1Scr (C) on HNE (25nM) protease activity, using three molar ratios. (D-E) Relative intensity of the various cleaved forms of HisBMG-1 and HisBMG-1Scr, incubated with HNE at (D) 2.5 : 1 and (E) 10 : 1 ratio; identified by native mass spectrometry. Peak intensities were normalised to the highest intensity signal emitted by the 1.8 kD peak. (F-G) Permeability of colonic epithelial cells (day 10-12 ALI) pretreated with 500nM HNE ± HisBMG-1 (F) or HisBMG-1Scr (G), over 6h time-course. (H) Colon epithelium permeability at 6h post HNE ± HisBMG-1 or HisBMG-1Scr treatment, extracted from (E) and (F). All data is presented as mean ± SEM, n=7 (A), n=4 (B), n=3 (C-H). p-adj<0.05 (*), p-adj<0.01 (**), Treatments were compared using repeated measures two-way ANOVA (A-C, F-G), multiple unpaired t-test (D-E); or one-way ANOVA (H). Mass spectrometry replicates are three independent measurements. Significance between HNE and HisBMG-1/HisBMG-1Scr is marked by an asterisk matching the dose colour.

To more precisely determine the interaction between BMG-1 and HNE, mass spectrometry analysis was performed of the HisBMG-1 and the control HisBMG-1Scr, preincubated with HNE using two different ratios of the microbial protein and HNE (2.5:1 and 10:1) (Figure 5D-E). While HisBMG-1 and HisBMG-1Scr alone displayed the intact protein spectra only (Figure S5-D and G), incubation with HNE resulted in 4 different products: an intact protein and 3 cleaved products with 1.2 kDa, 1.8 kDa and 2.7 kDa loss in mass (Figure S5-E, S5-F, S5-H and S5-I). Following HNE incubation, the abundance of the intact product was significantly higher in HisBMG-1Scr control compared to HisBMG-1 (Figure 5E). In contrast, 1.2 and 2.7 kDa products demonstrated significantly higher abundance in HisBMG-1 interactions with HNE, indicating active cleavage (Figure 5D-E). These data demonstrate a significantly higher interaction, binding affinity, and cleavage process of HNE for HisBMG-1 compared to the scrambled control, HisBMG-1Scr. Finally, we evaluated the protective effect of HisBMG-1 on differentiated colon epithelial cells specifically against HNE exposure (rather than total neutrophil contents), and demonstrated a dose-dependent protective effect, while HisBMG-1Scr failed to show this effect (Figure 5F-H). These data demonstrates a high specificity of HisBMG-1 towards the human neutrophil elastase, a key protease with known barrier damaging properties in the colonic inflammation.

### BMG-1 promotes colonic epithelial wound healing and prevents colon epithelial cell death

Given BMG-1 protected the colonic epithelial barrier integrity from increased permeability, we next examined its capacity to promote colon epithelial repair after damage, a common pathology in IBD. We performed a 2-dimensional colonic epithelial scratch wound assay using human colonic organoid-derived cells, by introducing the wound and incubating the cells with HNE premixed with HisBMG-1 or HisBMG-1Scr. HNE alone significantly delayed wound healing by more than 50% over baseline, but cells treated with HisBMG-1 exhibited accelerated healing and 100% complete healing at all doses of HisBMG-1 tested (Figure 6A). This effect was slightly stronger than that of elafin (Figure S6A-C). In contrast, HisBMG-1Scr failed to have this potent effect on wound healing at low doses, however, at higher doses it was able to achieve this a protective effect suggesting that unlike barrier permeability the impact of BMG-1 on wound healing is not completely due to the anti-protease RCL domain (Figure 6B and S6C). Healing of the colonic barrier was tested with HNE at 100 nM, therefore next we aimed to test whether HisBMG-1 could impact colonic epithelial cell death when exposed to a higher dose of HNE (250 nM) that could induce clear cell death. We exposed the colon epithelial cells to HNE and increasing doses of HisBMG-1 for 24h, measuring cell confluency and metabolic activity (MTT). HisBMG-1 at 2uM, 1uM and 0.5uM, fully protected against cell death and loss of confluency caused by high dose HNE (Figure 6C and S6-D), which was completely absent in HisBMG-1Scr treated cells (Figure 6D and S6D). These results were confirmed with the MTT assay, where HisBMG-1 preserved cell viability in a dose-dependent manner and this was again lost with HisBMG-1Scr (Figure 6E-F). These results clearly demonstrate a strong and specific protective effect of BMG-1 on epithelial repair and viability.

**Figure 6.**
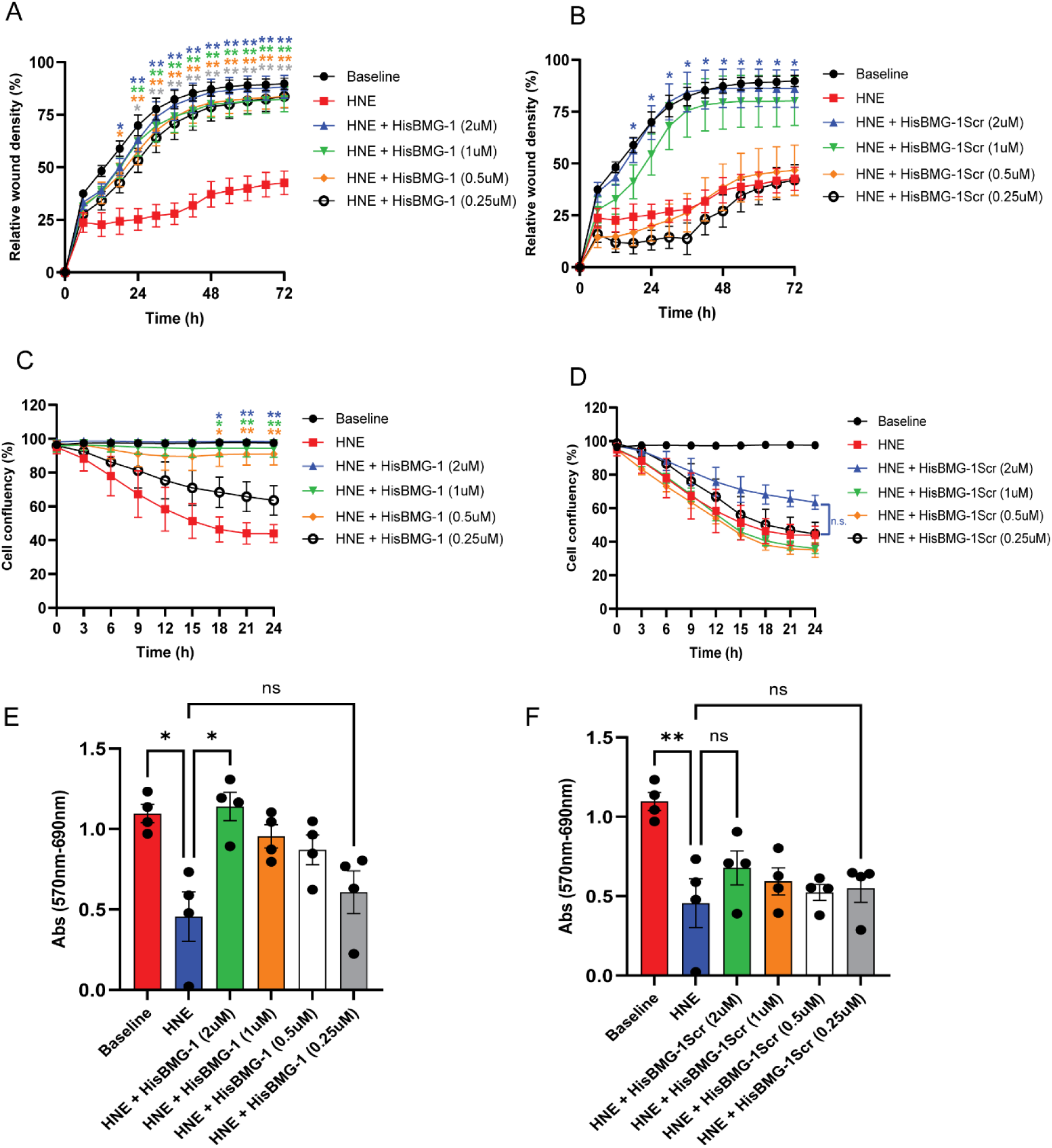
HisBMG-1 promotes colon epithelial barrier healing and prevents human neutrophil elastase-induced cell death. (A-B) The effect of HisBMG-1 (A) and HisBMG-1Scr (B) on relative wound density in colon epithelial cells treated with 100nM HNE ± HisBMG-1/HisBMG-1Scr healing for 72h post-wounding. (C-D) The effect of HisBMG-1 (C) and HisBMG-1Scr (D) on cell layer confluency in colon epithelial cells treated with 250nM HNE ± HisBMG-1/HisBMG-1Scr for 24h. (E-F) Colon epithelial cell viability after 24h treatment with 250nM HNE ± HisBMG-1 (E) or HisBMG-1Scr (F), measured by MTT assay. All data is presented as mean ± SEM, n=7 (A), n=4 (B-F). p-adj<0.05 (*), p-adj<0.01 (**), Treatments were compared using repeated measures two-way ANOVA (A-D); or Friedman test (E-F). Significance between HNE and HisBMG-1/HisBMG-1Scr is marked by an asterisk matching the dose colour.

### BMG-1 reduces colon damage in DSS-induced colitis in vivo

After demonstrating the capacity of HisBMG-1 for HNE inhibition, and colon epithelium protection *in vitro*, we aimed to test its capacity to alter the disease course in a murine model of DSS colitis *in vivo*. BMG-1 specifically targets human neutrophil elastase, which only shares 71% sequence similarity and identity with mouse neutrophil elastase. Despite this, BMG-1 is still able to partially inhibit the activity of mouse neutrophil proteases (∼30% reduction in protease activity, data not shown), albeit to a far lesser extent than human neutrophil proteases. Mice were treated with 2% DSS in drinking water, followed by recovery with normal drinking water. HisBMG-1 was administrated intra-rectally every two days (Figure 7A). We did not observe differences in weight loss between DSS mice treated with vehicle or HisBMG-1 (Figure 7B). DSS treatment induced a significant increase in the protease activity within the intestinal tract as evidenced by increased faecal protease activity compared to control drinking water groups (Figure 7C-D). HisBMG-1 administration at least partly abrogated this elevation in protease activity above non-drinking water controls (Figure 7C-D). As expected, the DSS-induced mucosal damage and ulceration led to a significant loss of histologically normal colon in DSS-treated mice and a significantly shorter colon length than control drinking water mice (Figure 7E-G). This colonic damage was reduced in HisBMG-1 treated mice as compared to the vehicle controls resulting in a significantly greater percentage of normal colon tissue (i.e. less damaged tissue) and greater colon length (Figure 7E-G).

**Figure 7.**
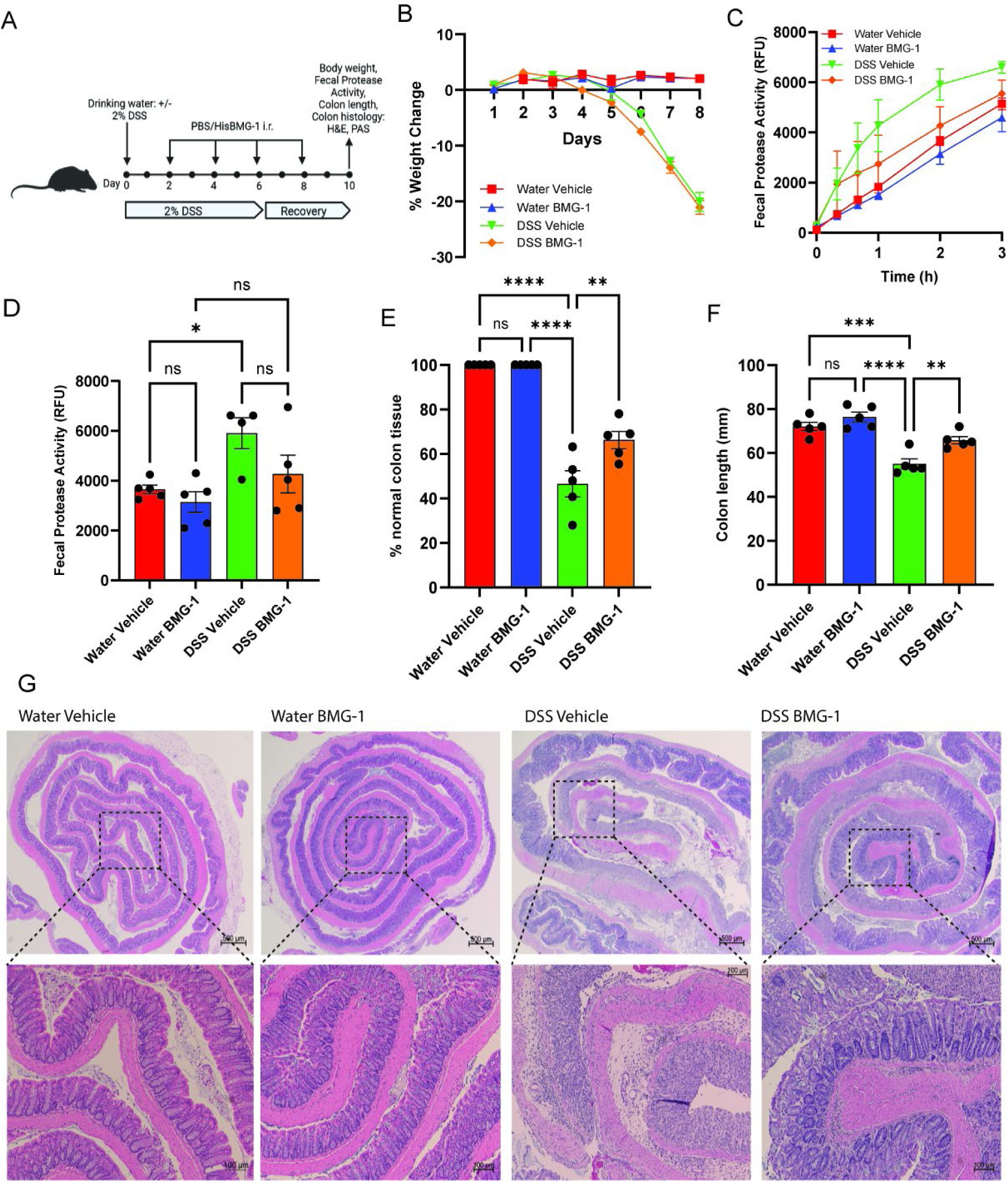
Administration of HisBMG-1 reduces DSS-induced colon damage in mice. (A) Schematic overview of the experimental design. i.r. – intrarectal, endpoint day 8 DSS, and day 10 drinking water. (B) Body weight change. (C) Faecal protease activity measured over 3h, in endpoint samples, using a broad spectrum protease substrate. (D) Faecal protease activity at 2h, extracted from (C). (E) Colon histopathology assessment by H&E PAS at endpoint. (F) Macroscopic colon length at endpoint. (G) Representative images of H&E stained colon sections in four experimental groups. Top row (250x magnification), bottom row (1000x magnification). All data is presented as mean ± SEM, n=4-5. p-adj<0.05 (*), p-adj<0.01 (**), p-adj<0.001 (***), p-adj<0.0001 (****), Treatments were compared using ordinary one-way ANOVA and Dunnett’s multiple comparison test.

## Discussion

Through the design of a new systematic intestinal microbiome metatranscriptomic pipeline and functional screening platform, we identified, synthesised and functionally tested BMG-1, a novel and potent neutrophil elastase inhibitor, with colonic epithelial protective properties. This previously uncharacterised microbial protein showed gene expression that inversely correlated with disease activity, that is, the higher the activity (disease flares) the less expression of the gene. BMG-1 inhibits human neutrophil elastase and neutrophil cell proteolytic activity. Importantly, BMG-1 protects the colonic epithelial barrier from proteolytic damage, promotes colon epithelial wound healing and reduces colon damage in a mouse model of colitis.

Previous studies using bioinformatics to investigate the microbiome in the context of IBD have mainly focussed on either taxonomy, metabolites, or broad categories of functional units of genes such as KEGG categories ^52,53^. These studies for the most part do not investigate specific gene/protein features and their functionality in disease, nor how specific gene features are linked to disease activity and *in silico* secretion prediction models. To develop novel therapeutic treatments to more effectively tackle unmet clinical needs we aimed to leverage the microbiome’s role in healthy gut homeostasis, and hypothesised that identifying functional microbial products through transcriptional interrogation would provide novel candidates. Our pipeline was specifically designed to prioritize the activity and functionality of the microbiota by incorporating microbial metatranscriptomic analysis of stool samples to identify actively transcribed protein-coding RNA.

A fundamental aspect of this bioinformatic based approach involved the construction of personalized genomic references (PGRs), followed by transcriptome mapping to these references. Notably, we observed that ∼10-15% of total reads remained unmapped to the reference. This incompleteness was anticipated and arises from several factors, including the abundance of uncharacterised species within the human gut metagenome, and the variable degree of completeness of the reference genomes used for mapping. Specifically, the latter has been a point of discussion among researchers, and it is widely accepted that approximately one third of the microbial genes remain functionally uncharacterised ^54 55^. These factors primarily reflect the inherent challenges and developmental stage of the evolving field of functionality within microbiome research.

Most bioactive microbiota molecules linked to disease or healthy states as well as microbiota-derived drug candidates have been metabolite-based. The utility of protein-based secreted microbial therapeutics has remained largely unexplored. Our study is among the first to identify novel functional microbiota-derived secreted proteins with therapeutic potential for IBD through the meta-transcriptomic interrogation of stool samples. We utilised bioassays that were designed to assess the impact on hallmark features of IBD-related pathology, including impaired epithelial integrity and wound repair, for functional validation of our identified microbial protein targets. In our assays we employed neutrophils to damage the barrier. Elevated neutrophil-derived molecules, such as neutrophil elastase, are abundant in the lumen and strongly associated with IBD diagnosis and disease activity ^14,56,57^. Our findings suggest that BMG-1 may exert its protective effect by inhibiting neutrophil elastase at the epithelial surface preventing epithelial damage and subsequent barrier leakage.

Dysregulated protease activity in the gastrointestinal (GI) tract is being increasingly linked to IBD, with elevated protease levels observed in patient faecal samples, even preceding diagnosis ^14,58–61^. This imbalance can arise from either increased proteolytic enzymes, or due to the impaired inhibition of these enzymes. These enzymes are largely either digestive enzymes derived from the pancreas or produced by immune cells such as mast cells and neutrophils ^58^. Importantly, proteases derived from neutrophils, such as HNE and MMPs, are strongly correlated with IBD severity ^62^. Upon inflammatory insult, neutrophils are recruited to eliminate bacteria through phagocytosis, NETs, and ROS production. Apart from ROS, these responses involve the release of proteolytic enzymes that as a side effect of targeting microbes also degrade tight junction proteins and compromise barrier integrity. Inhibiting HNE may reduce elastolytic activity of granules and NET-associated enzymes which may potentially limit the leakiness of barrier function in IBD.

Recent studies suggest that the gut microbiota may also directly produce detrimental proteases, exacerbating IBD pathology ^13,14^. Certain bacterial species, such as *P. vulgatus* and *E. faecalis*, produce proteases that have been linked to disease activity ^13,63^. However, an alternative hypothesis suggests that the absence of protease inhibitors, rather than excessive protease production, drives colonic damage and inflammation ^64^. In homeostasis, the gut maintains protease balance through multiple host inhibitors, including alpha1-antitrypsin (AAT), SLPI, and Elafin. While AAT primarily functions in serum, SLPI and Elafin neutralize neutrophil-derived proteases in the gut mucosa ^65,66^. Notably, Elafin and SLPI expression are downregulated in mucosal biopsies of IBD patients, potentially compromising epithelial barrier protection ^22,67^. Studies have also demonstrated that protease inhibition in the gut can alleviate inflammation in mouse models of colitis ^68–70^. These studies suggest that during chronic inflammation, the epithelial barrier has a dysfunctional host anti-protease/proteolytic balance. However, the focus of these studies has been on synthetic or host-derived protease inhibitors, ignoring the role of microbiota protease inhibition. Our study is one of the first to demonstrate that the gut microbiome can also play a key role to suppress protease activity in IBD. Identifying a specific microbial protein BMG-1 that acts as a potent inhibitor of HNE. BMG-1’s expression correlates with disease remission and flares, thereby suggesting a role for maintaining the proteolytic balance in the gut. Hence, BMG-1 may also represent a candidate for future therapeutic targeting, potentially enhancing epithelial barrier integrity and repair in IBD. Only four proteins with protease inhibitory properties have been functionally characterised in gut bacteria to date ^71^, isolated from *Bifidobacterium longum* ^50^, *Eubacterium sireaum* ^43^ and *Eubacterium saburreum* ^49^. However, none of these proteins have been investigated in the context of a disease, including no link to IBD. We identified the bacterium that likely expresses the BMG-1 gene in the gut as *F. plautii*, known as an anaerobic gram-positive member of the healthy human gut microbiome. BMG-1 naturally produced by *F. plautii* may neutralise HNE activity in order to perform a symbiotic role for both the host and bacteria to help maintain mucosal barrier integrity and hence maintain the homeostatic niche of this mucosal microbe. The role of *F. plautii* in IBD has remained relatively unexplored with only a few studies describing its effects in mouse models ^72,73^.

To our knowledge, BMG-1 is the first microbiota-derived protein drug candidate generated through transcriptomic gene expression analysis of the gut microbiome. Our study suggests that the balance of protease activity regulated by both host and microbial factors could be involved as a potential factor in IBD pathogenesis. Our data reveals that the inhibition of HNE by BMG-1 may be a part of the mechanism by which the microbiome contributes to the balance of the gut protease/anti-protease homeostasis. BMG-1 represents a novel candidate for future therapeutic targeting, potentially enhancing epithelial barrier integrity and repair in IBD.

## Supplementary figure legends

**Figure S1.** Summary of quality control steps and gene filtering procedures in the metatranscriptomic analysis of the gut microbiome in IBD. (A) Stacked column chart representing overall percentage of read mapping to personalised (PGR) and global genomic references (GGR). (B) Multiple sequence alignment of 12 representative genes, clustered based on 98% shared amino acid sequence identity. Coding sequences could differ 2% in length and identity. (C) Faecal Calprotectin (FC) levels of stool samples used in disease activity analysis stratified by high and low disease activity based on 200 µg/g stool cutoff. (D, E) Number of gene coding features that are detected in a given fraction of stool metatranscriptomic sample timepoints. Threshold for feature detection in a sample is > 5 reads. (F, G) Venn Diagram on gene lists from 4 groups; Two statistical models (VarPart and DESeq2) and two clustering strategies (98 and 100%) used for both disease group comparisons, CD vs non-IBD (F) and UC vs non-IBD (G). (H, I, J) Pie charts representing Enzyme Classification System (EC system) categories from top DE genes in CD vs non-IBD (752 genes), UC vs non-IBD (2134 genes), and high vs low disease activity (1177 genes).

**Figure S2.** Taxonomic dot plot of taxa predicted to express the top differentially expressed genes. Species expressing genes with Log2FC > 2 and p-value < 10^-4^ (∼9000 unique genes) were included in the analysis. Phyla, family, genus and species is used as taxa ordering criterion. Vertical columns represent different disease groups comparisons and cluster strategies (98% and 100% amino acid sequence identity similarity). Dot colour indicates p-value of genes, with the minimum p-value used if multiple genes come from the same species. Dot size represents the percentage of samples in which the gene is present (or averaged percentage if multiple genes are expressed by a single species).

**Figure S3.** Biomarker analysis using Receiver Operating Characteristic (ROC) curves. Two analytical approaches were employed: (1) pairwise combinations of gene features, and (2) Random Forest classification using 500 trees, using DE genes from the DESeq2 analysis with an adjusted p-value < 1×10⁻⁵ for each contrast group. (A, B, C) ROC curve of top result from pairwise feature combinations for each disease group comparison, CD vs non-IBD (A), UC vs non-IBD (B) and high vs low disease activity (C). (D, E, F) ROC curves of top predictive features obtained using a random forest model, for each disease group comparison, CD vs non-IBD (D), UC vs non-IBD (E) and high vs low disease activity (F). (G, H, I) Mean Decrease Accuracy and Mean Decrease Gini plots for each disease group comparison, CD vs non-IBD (G), UC vs non-IBD (H) and high vs low disease activity (I).

**Figure S4.** BMG-1 inhibits neutrophil protease activity and protects the colon epithelium from damage. (A-B) Barrier permeability colon epithelial cells (air-liquid interface day 2-4) pretreated with neutrophil lysate (NeutLys) ± BMG-1 (A) or Elafin (B), measured over 6h time-course. (C) The protein sequence alignment between BMG-1 and BMG-2, showing 13 amino acid substitutions highlighted by red arrows. (D-E) The effect of BMG-1 (D) or Elafin (E) on neutrophil lysate protease activity in a cell-free assay. (F) Scrambling mutations of the RCL protein sequence in four HisBMG-1Scr candidates (left) and corresponding SDS-PAGE expression profile showing bacterial lysate (L) and purified protein (P) for each of the candidates. Four irrelevant lanes were cropped out from the right-hand side of the gel image. (G) Human Neutrophil Elastase protease inhibition by HisBMG-1Scr candidates, using four HNE : HisBMG-1Scr molar ratios. HisBMG-1Scr2 was selected for all downstream experiments. All data is presented as mean ± SEM; n=6 (A), n=5 (B), n=4 (D), n=3 (E), n=1 (F-G mean ± SD). p-adj<0.05 (*), p-adj<0.01 (**), Treatments were compared using repeated measures two-way ANOVA and Dunnett’s multiple comparison test. Significance between NeutLys and HisBMG-1/HisBMG-1Scr is marked by an asterisk matching the dose colour.

**Figure S5.** BMG-1 is a highly specific inhibitor of human neutrophil elastase (HNE). (A-C) The effect of BMG-1 on HNE (25nM) (A), Cathepsin G (50nM) (B) and Proteinase 3 (50nM) (C) protease activity, using three molar ratios between the enzyme and the inhibitor. (D-I) Native mass spectra: HisBMG-1 (D) HNE : HisBMG-1 (1 : 2.5) (E), HNE : HisBMG-1 (1 : 10) (F), HisBMG-1Scr (G), HNE : HisBMG-1Scr (1 : 2.5) (H), HNE : HisBMG-1Scr (1 : 10) (I). The data in A is presented as mean ± SEM, n=4, data in B and C represents a single representative experiment. p-adj<0.05 (*), p-adj<0.01 (**), Treatments were compared using repeated measures two-way ANOVA (A). Significance between HNE and BMG-1 is marked by an asterisk matching the dose colour.

**Figure S6.** BMG-1 promotes colon epithelial healing and prevents HNE-induced cell death. (A-B) The effect of BMG-1 (A) and Elafin (B) on relative wound density in colon epithelial cells treated with 100nM HNE ± BMG-1/Elafin for 72h of healing post-scratch wounding. (C-D) Representative phase contrast images of colon epithelial cells treated with 100nM HNE ± HisBMG-1/HisBMG-1Scr for 72h post-scratch wounding (C) and after 24h 250nM HNE ± HisBMG-1/HisBMG-1Scr (D). Contrast was increased to 84% on all images in C and D, to visualise clear wound or cell death boundaries. All data is presented as mean ± SEM, n=7 (A-B). p-adj<0.05 (*), p-adj<0.01 (**). Treatments were compared using repeated measures two-way ANOVA. Significance between HNE and BMG-1/Elafin is marked by an asterisk matching the dose colour.

## Acknowledgements

funding for this study was provided by McCusker Charitable Foundation, National Health and Medical Research Council of Australia, as well as internal University of Newcastle research funding.

## References

1. Kang, L., Fang, X., Song, Y.H., He, Z.X., Wang, Z.J., Wang, S.L., Li, Z.S., and Bai, Y. (2022). Neutrophil-Epithelial Crosstalk During Intestinal Inflammation. Cell Mol Gastroenterol Hepatol 14, 1257–1267. 10.1016/j.jcmgh.2022.09.002.

2. Le Berre, C., Honap, S., and Peyrin-Biroulet, L. (2023). Ulcerative colitis. Lancet 402, 571–584. 10.1016/S0140-6736(23)00966-2.

3. Torres, J., Mehandru, S., Colombel, J.F., and Peyrin-Biroulet, L. (2017). Crohn’s disease. Lancet 389, 1741–1755. 10.1016/S0140-6736(16)31711-1.

4. Gibble, T.H., Naegeli, A.N., Grabner, M., Isenberg, K., Shan, M., Teng, C.C., and Curtis, J.R. (2023). Identification of inadequate responders to advanced therapy among commercially-insured adult patients with Crohn’s disease and ulcerative colitis in the United States. BMC Gastroenterol 23, 63. 10.1186/s12876-023-02675-w.

5. Roda, G., Jharap, B., Neeraj, N., and Colombel, J.F. (2016). Loss of Response to Anti-TNFs: Definition, Epidemiology, and Management. Clin Transl Gastroenterol 7, e135. 10.1038/ctg.2015.63.

6. Irvine, E.J., and Marshall, J.K. (2000). Increased intestinal permeability precedes the onset of Crohn’s disease in a subject with familial risk. Gastroenterology 119, 1740–1744. 10.1053/gast.2000.20231.

7. Turpin, W., Lee, S.H., Raygoza Garay, J.A., Madsen, K.L., Meddings, J.B., Bedrani, L., Power, N., Espin-Garcia, O., Xu, W., Smith, M.I., et al. (2020). Increased Intestinal Permeability Is Associated With Later Development of Crohn’s Disease. Gastroenterology 159, 2092–2100 e2095. 10.1053/j.gastro.2020.08.005.

8. Parkes, G., Ungaro, R.C., Danese, S., Abreu, M.T., Arenson, E., Zhou, W., Ilo, D., Laroux, F.S., Deng, H., Sanchez Gonzalez, Y., and Peyrin-Biroulet, L. (2023). Correlation of mucosal healing endpoints with long-term clinical and patient-reported outcomes in ulcerative colitis. J Gastroenterol 58, 990–1002. 10.1007/s00535-023-02013-7.

9. Shah, S.C., Colombel, J.F., Sands, B.E., and Narula, N. (2016). Mucosal Healing Is Associated With Improved Long-term Outcomes of Patients With Ulcerative Colitis: A Systematic Review and Meta-analysis. Clin Gastroenterol Hepatol 14, 1245–1255 e1248. 10.1016/j.cgh.2016.01.015.

10. Bressenot, A., Salleron, J., Bastien, C., Danese, S., Boulagnon-Rombi, C., and Peyrin-Biroulet, L. (2015). Comparing histological activity indexes in UC. Gut 64, 1412–1418. 10.1136/gutjnl-2014-307477.

11. Danne, C., Skerniskyte, J., Marteyn, B., and Sokol, H. (2024). Neutrophils: from IBD to the gut microbiota. Nat Rev Gastroenterol Hepatol 21, 184–197. 10.1038/s41575-023-00871-3.

12. Jablaoui, A., Kriaa, A., Mkaouar, H., Akermi, N., Soussou, S., Wysocka, M., Woloszyn, D., Amouri, A., Gargouri, A., Maguin, E., et al. (2020). Fecal Serine Protease Profiling in Inflammatory Bowel Diseases. Front Cell Infect Microbiol 10, 21. 10.3389/fcimb.2020.00021.

13. Mills, R.H., Dulai, P.S., Vazquez-Baeza, Y., Sauceda, C., Daniel, N., Gerner, R.R., Batachari, L.E., Malfavon, M., Zhu, Q., Weldon, K., et al. (2022). Multi-omics analyses of the ulcerative colitis gut microbiome link Bacteroides vulgatus proteases with disease severity. Nat Microbiol 7, 262–276. 10.1038/s41564-021-01050-3.

14. Galipeau, H.J., Caminero, A., Turpin, W., Bermudez-Brito, M., Santiago, A., Libertucci, J., Constante, M., Raygoza Garay, J.A., Rueda, G., Armstrong, S., et al. (2021). Novel Fecal Biomarkers That Precede Clinical Diagnosis of Ulcerative Colitis. Gastroenterology 160, 1532–1545. 10.1053/j.gastro.2020.12.004.

15. Manichanh, C., Rigottier-Gois, L., Bonnaud, E., Gloux, K., Pelletier, E., Frangeul, L., Nalin, R., Jarrin, C., Chardon, P., Marteau, P., et al. (2006). Reduced diversity of faecal microbiota in Crohn’s disease revealed by a metagenomic approach. Gut 55, 205–211. 10.1136/gut.2005.073817.

16. Ott, S.J., Musfeldt, M., Wenderoth, D.F., Hampe, J., Brant, O., Folsch, U.R., Timmis, K.N., and Schreiber, S. (2004). Reduction in diversity of the colonic mucosa associated bacterial microflora in patients with active inflammatory bowel disease. Gut 53, 685–693. 10.1136/gut.2003.025403.

17. Haifer, C., Paramsothy, S., Kaakoush, N.O., Saikal, A., Ghaly, S., Yang, T., Luu, L.D.W., Borody, T.J., and Leong, R.W. (2022). Lyophilised oral faecal microbiota transplantation for ulcerative colitis (LOTUS): a randomised, double-blind, placebo-controlled trial. Lancet Gastroenterol Hepatol 7, 141–151. 10.1016/S2468-1253(21)00400-3.

18. Moayyedi, P., Surette, M.G., Kim, P.T., Libertucci, J., Wolfe, M., Onischi, C., Armstrong, D., Marshall, J.K., Kassam, Z., Reinisch, W., and Lee, C.H. (2015). Fecal Microbiota Transplantation Induces Remission in Patients With Active Ulcerative Colitis in a Randomized Controlled Trial. Gastroenterology 149, 102–109 e106. 10.1053/j.gastro.2015.04.001.

19. Paramsothy, S., Kamm, M.A., Kaakoush, N.O., Walsh, A.J., van den Bogaerde, J., Samuel, D., Leong, R.W.L., Connor, S., Ng, W., Paramsothy, R., et al. (2017). Multidonor intensive faecal microbiota transplantation for active ulcerative colitis: a randomised placebo-controlled trial. Lancet 389, 1218–1228. 10.1016/S0140-6736(17)30182-4.

20. Feng, J., Chen, Y., Liu, Y., Lin, L., Lin, X., Gong, W., Xia, R., He, J., Sheng, J., Cai, H., and Xiao, C. (2023). Efficacy and safety of fecal microbiota transplantation in the treatment of ulcerative colitis: a systematic review and meta-analysis. Sci Rep 13, 14494. 10.1038/s41598-023-41182-6.

21. Morohoshi, Y., Matsuoka, K., Chinen, H., Kamada, N., Sato, T., Hisamatsu, T., Okamoto, S., Inoue, N., Takaishi, H., Ogata, H., et al. (2006). Inhibition of neutrophil elastase prevents the development of murine dextran sulfate sodium-induced colitis. J Gastroenterol 41, 318–324. 10.1007/s00535-005-1768-8.

22. Motta, J.P., Bermudez-Humaran, L.G., Deraison, C., Martin, L., Rolland, C., Rousset, P., Boue, J., Dietrich, G., Chapman, K., Kharrat, P., et al. (2012). Food-grade bacteria expressing elafin protect against inflammation and restore colon homeostasis. Sci Transl Med 4, 158ra144. 10.1126/scitranslmed.3004212.

23. Integrative, H.M.P.R.N.C. (2019). The Integrative Human Microbiome Project. Nature 569, 641–648. 10.1038/s41586-019-1238-8.

24. Lloyd-Price, J., Arze, C., Ananthakrishnan, A.N., Schirmer, M., Avila-Pacheco, J., Poon, T.W., Andrews, E., Ajami, N.J., Bonham, K.S., Brislawn, C.J., et al. (2019). Multi-omics of the gut microbial ecosystem in inflammatory bowel diseases. Nature 569, 655–662. 10.1038/s41586-019-1237-9.

25. Truong, D.T., Franzosa, E.A., Tickle, T.L., Scholz, M., Weingart, G., Pasolli, E., Tett, A., Huttenhower, C., and Segata, N. (2015). MetaPhlAn2 for enhanced metagenomic taxonomic profiling. Nat Methods 12, 902–903. 10.1038/nmeth.3589.

26. Langmead, B., and Salzberg, S.L. (2012). Fast gapped-read alignment with Bowtie 2. Nat Methods 9, 357–359. 10.1038/nmeth.1923.

27. Seemann, T. (2014). Prokka: rapid prokaryotic genome annotation. Bioinformatics 30, 2068–2069. 10.1093/bioinformatics/btu153.

28. Lorenz, R., Bernhart, S.H., Honer Zu Siederdissen, C., Tafer, H., Flamm, C., Stadler, P.F., and Hofacker, I.L. (2011). ViennaRNA Package 2.0. Algorithms Mol Biol 6, 26. 10.1186/1748-7188-6-26.

29. Fu, L., Niu, B., Zhu, Z., Wu, S., and Li, W. (2012). CD-HIT: accelerated for clustering the next-generation sequencing data. Bioinformatics 28, 3150–3152. 10.1093/bioinformatics/bts565.

30. Liao, Y., Smyth, G.K., and Shi, W. (2013). The Subread aligner: fast, accurate and scalable read mapping by seed-and-vote. Nucleic Acids Res 41, e108. 10.1093/nar/gkt214.

31. Lamb, C.A., Kennedy, N.A., Raine, T., Hendy, P.A., Smith, P.J., Limdi, J.K., Hayee, B., Lomer, M.C.E., Parkes, G.C., Selinger, C., et al. (2019). British Society of Gastroenterology consensus guidelines on the management of inflammatory bowel disease in adults. Gut 68, s1–s106. 10.1136/gutjnl-2019-318484.

32. Hoffman, G.E., and Schadt, E.E. (2016). variancePartition: interpreting drivers of variation in complex gene expression studies. BMC Bioinformatics 17, 483. 10.1186/s12859-016-1323-z.

33. Love, M.I., Huber, W., and Anders, S. (2014). Moderated estimation of fold change and dispersion for RNA-seq data with DESeq2. Genome Biol 15, 550. 10.1186/s13059-014-0550-8.

34. Ritchie, M.E., Phipson, B., Wu, D., Hu, Y., Law, C.W., Shi, W., and Smyth, G.K. (2015). limma powers differential expression analyses for RNA-sequencing and microarray studies. Nucleic Acids Res 43, e47. 10.1093/nar/gkv007.

35. Almagro Armenteros, J.J., Tsirigos, K.D., Sonderby, C.K., Petersen, T.N., Winther, O., Brunak, S., von Heijne, G., and Nielsen, H. (2019). SignalP 5.0 improves signal peptide predictions using deep neural networks. Nat Biotechnol 37, 420–423. 10.1038/s41587-019-0036-z.

36. Eichinger, V., Nussbaumer, T., Platzer, A., Jehl, M.A., Arnold, R., and Rattei, T. (2016). EffectiveDB--updates and novel features for a better annotation of bacterial secreted proteins and Type III, IV, VI secretion systems. Nucleic Acids Res 44, D669–674. 10.1093/nar/gkv1269.

37. Huang, X., Kamadurai, H., Siuti, P., Ahmed, E., Bennett, J.L., and Donald, W.A. (2023). Oligomeric Remodeling by Molecular Glues Revealed Using Native Mass Spectrometry and Mass Photometry. J Am Chem Soc 145, 14716–14726. 10.1021/jacs.3c02655.

38. Bruce, J., Kaiko, G.E., and Keely, S. (2019). Isolation and In Vitro Culture of Human Gut Progenitor Cells. Methods Mol Biol 2029, 49–62. 10.1007/978-1-4939-9631-5_5.

39. Kaiko, G.E., Ryu, S.H., Koues, O.I., Collins, P.L., Solnica-Krezel, L., Pearce, E.J., Pearce, E.L., Oltz, E.M., and Stappenbeck, T.S. (2016). The Colonic Crypt Protects Stem Cells from Microbiota-Derived Metabolites. Cell 167, 1137. 10.1016/j.cell.2016.10.034.

40. Miyoshi, H., Ajima, R., Luo, C.T., Yamaguchi, T.P., and Stappenbeck, T.S. (2012). Wnt5a potentiates TGF-beta signaling to promote colonic crypt regeneration after tissue injury. Science 338, 108–113. 10.1126/science.1223821.

41. Miyoshi, H., and Stappenbeck, T.S. (2013). In vitro expansion and genetic modification of gastrointestinal stem cells in spheroid culture. Nat Protoc 8, 2471–2482. 10.1038/nprot.2013.153.

42. VanDussen, K.L., Marinshaw, J.M., Shaikh, N., Miyoshi, H., Moon, C., Tarr, P.I., Ciorba, M.A., and Stappenbeck, T.S. (2015). Development of an enhanced human gastrointestinal epithelial culture system to facilitate patient-based assays. Gut 64, 911–920. 10.1136/gutjnl-2013-306651.

43. Mkaouar, H., Akermi, N., Mariaule, V., Boudebbouze, S., Gaci, N., Szukala, F., Pons, N., Marquez, J., Gargouri, A., Maguin, E., and Rhimi, M. (2016). Siropins, novel serine protease inhibitors from gut microbiota acting on human proteases involved in inflammatory bowel diseases. Microb Cell Fact 15, 201. 10.1186/s12934-016-0596-2.

44. Kaiko, G.E., Chen, F., Lai, C.W., Chiang, I.L., Perrigoue, J., Stojmirovic, A., Li, K., Muegge, B.D., Jain, U., VanDussen, K.L., et al. (2019). PAI-1 augments mucosal damage in colitis. Sci Transl Med 11. 10.1126/scitranslmed.aat0852.

45. Ananthakrishnan, A.N., Luo, C., Yajnik, V., Khalili, H., Garber, J.J., Stevens, B.W., Cleland, T., and Xavier, R.J. (2017). Gut microbiome function predicts response to anti-integrin biologic therapy in inflammatory bowel diseases. Cell host & microbe 21, 603–610. e603.

46. Cortez, R.V., Moreira, L.N., Padilha, M., Bibas, M.D., Toma, R.K., Porta, G., and Taddei, C.R. (2021). Gut microbiome of children and adolescents with primary sclerosing cholangitis in association with ulcerative colitis. Frontiers in Immunology 11, 598152.

47. Machiels, K., Joossens, M., Sabino, J., De Preter, V., Arijs, I., Eeckhaut, V., Ballet, V., Claes, K., Van Immerseel, F., and Verbeke, K. (2014). A decrease of the butyrate-producing species Roseburia hominis and Faecalibacterium prausnitzii defines dysbiosis in patients with ulcerative colitis. Gut 63, 1275–1283.

48. Sokol, H., Pigneur, B., Watterlot, L., Lakhdari, O., Bermúdez-Humarán, L.G., Gratadoux, J.-J., Blugeon, S., Bridonneau, C., Furet, J.-P., and Corthier, G. (2008). Faecalibacterium prausnitzii is an anti-inflammatory commensal bacterium identified by gut microbiota analysis of Crohn disease patients. Proceedings of the National Academy of Sciences 105, 16731–16736.

49. Akermi, N., Mkaouar, H., Kriaa, A., Jablaoui, A., Soussou, S., Gargouri, A., Coleman, A.W., Perret, F., Maguin, E., and Rhimi, M. (2019). para-Sulphonato-calix[n]arene capped silver nanoparticles challenge the catalytic efficiency and the stability of a novel human gut serine protease inhibitor. Chem Commun (Camb) 55, 8935–8938. 10.1039/c9cc03183a.

50. Ivanov, D., Emonet, C., Foata, F., Affolter, M., Delley, M., Fisseha, M., Blum-Sperisen, S., Kochhar, S., and Arigoni, F. (2006). A serpin from the gut bacterium Bifidobacterium longum inhibits eukaryotic elastase-like serine proteases. J Biol Chem 281, 17246–17252. 10.1074/jbc.M601678200.

51. Abramson, J., Adler, J., Dunger, J., Evans, R., Green, T., Pritzel, A., Ronneberger, O., Willmore, L., Ballard, A.J., Bambrick, J., et al. (2024). Accurate structure prediction of biomolecular interactions with AlphaFold 3. Nature 630, 493–500. 10.1038/s41586-024-07487-w.

52. Morgan, X.C., Tickle, T.L., Sokol, H., Gevers, D., Devaney, K.L., Ward, D.V., Reyes, J.A., Shah, S.A., LeLeiko, N., and Snapper, S.B. (2012). Dysfunction of the intestinal microbiome in inflammatory bowel disease and treatment. Genome biology 13, 1–18.

53. Gevers, D., Kugathasan, S., Denson, L.A., Vázquez-Baeza, Y., Van Treuren, W., Ren, B., Schwager, E., Knights, D., Song, S.J., and Yassour, M. (2014). The treatment-naive microbiome in new-onset Crohn’s disease. Cell host & microbe 15, 382–392.

54. Galperin, M.Y., and Koonin, E.V. (2004). ‘Conserved hypothetical’proteins: prioritization of targets for experimental study. Nucleic acids research 32, 5452–5463.

55. Bork, P. (2000). Powers and pitfalls in sequence analysis: the 70% hurdle. Genome research 10, 398–400.

56. Bohra, A., Mohamed, G., Vasudevan, A., Lewis, D., Van Langenberg, D.R., and Segal, J.P. (2023). The utility of faecal calprotectin, lactoferrin and other faecal biomarkers in discriminating endoscopic activity in Crohn’s disease: a systematic review and meta-analysis. Biomedicines 11, 1408.

57. Langhorst, J., Elsenbruch, S., Koelzer, J., Rueffer, A., Michalsen, A., and Dobos, G.J. (2008). Noninvasive markers in the assessment of intestinal inflammation in inflammatory bowel diseases: performance of fecal lactoferrin, calprotectin, and PMN-elastase, CRP, and clinical indices. Official journal of the American College of Gastroenterology| ACG 103, 162–169.

58. Motta, J.P., Magne, L., Descamps, D., Rolland, C., Squarzoni–Dale, C., Rousset, P., Martin, L., Cenac, N., Balloy, V., and Huerre, M. (2011). Modifying the protease, antiprotease pattern by elafin overexpression protects mice from colitis. Gastroenterology 140, 1272–1282.

59. Cenac, N., Andrews, C.N., Holzhausen, M., Chapman, K., Cottrell, G., Andrade-Gordon, P., Steinhoff, M., Barbara, G., Beck, P., and Bunnett, N.W. (2007). Role for protease activity in visceral pain in irritable bowel syndrome. The Journal of clinical investigation 117, 636–647.

60. Denadai-Souza, A., Bonnart, C., Tapias, N.S., Marcellin, M., Gilmore, B., Alric, L., Bonnet, D., Burlet-Schiltz, O., Hollenberg, M.D., and Vergnolle, N. (2018). Functional proteomic profiling of secreted serine proteases in health and inflammatory bowel disease. Scientific reports 8, 7834.

61. Li, Y., Watanabe, E., Kawashima, Y., Plichta, D.R., Wang, Z., Ujike, M., Ang, Q.Y., Wu, R., Furuichi, M., and Takeshita, K. (2022). Identification of trypsin-degrading commensals in the large intestine. Nature 609, 582–589.

62. Lakatos, G., Hritz, I., Varga, M.Z., Juhász, M., Miheller, P., Cierny, G., Tulassay, Z., and Herszényi, L. (2012). The impact of matrix metalloproteinases and their tissue inhibitors in inflammatory bowel diseases. Digestive Diseases 30, 289–295.

63. Steck, N., Hoffmann, M., Sava, I.G., Kim, S.C., Hahne, H., Tonkonogy, S.L., Mair, K., Krueger, D., Pruteanu, M., Shanahan, F., et al. (2011). Enterococcus faecalis metalloprotease compromises epithelial barrier and contributes to intestinal inflammation. Gastroenterology 141, 959–971. 10.1053/j.gastro.2011.05.035.

64. Qin, X.F. (2002). Impaired inactivation of digestive proteases by deconjugated bilirubin: the possible mechanism for inflammatory bowel disease. Med Hypotheses 59, 159–163. 10.1016/s0306-9877(02)00243-8.

65. Nugteren, S., and Samsom, J.N. (2021). Secretory Leukocyte Protease Inhibitor (SLPI) in mucosal tissues: Protects against inflammation, but promotes cancer. Cytokine & Growth Factor Reviews 59, 22–35.

66. Wiedow, O., Schröder, J., Gregory, H., Young, J., and Christophers, E. (1990). Elafin: an elastase-specific inhibitor of human skin. Purification, characterization, and complete amino acid sequence. Journal of Biological Chemistry 265, 14791–14795.

67. Schmid, M., Fellermann, K., Fritz, P., Wiedow, O., Stange, E.F., and Wehkamp, J. (2007). Attenuated induction of epithelial and leukocyte serine antiproteases elafin and secretory leukocyte protease inhibitor in Crohn’s disease. Journal of Leucocyte Biology 81, 907–915.

68. Ozaka, S., Sonoda, A., Ariki, S., Kamiyama, N., Hidano, S., Sachi, N., Ito, K., Kudo, Y., Minata, M., Saechue, B., et al. (2021). Protease inhibitory activity of secretory leukocyte protease inhibitor ameliorates murine experimental colitis by protecting the intestinal epithelial barrier. Genes Cells 26, 807–822. 10.1111/gtc.12888.

69. Van Spaendonk, H., Ceuleers, H., Smet, A., Berg, M., Joossens, J., Van der Veken, P., Francque, S.M., Lambeir, A.M., De Man, J.G., De Meester, I., et al. (2021). The Effect of a Novel Serine Protease Inhibitor on Inflammation and Intestinal Permeability in a Murine Colitis Transfer Model. Front Pharmacol 12, 682065. 10.3389/fphar.2021.682065.

70. Zhang, C., He, A., Liu, S., He, Q., Luo, Y., He, Z., Chen, Y., Tao, A., and Yan, J. (2019). Inhibition of HtrA2 alleviated dextran sulfate sodium (DSS)-induced colitis by preventing necroptosis of intestinal epithelial cells. Cell Death Dis 10, 344. 10.1038/s41419-019-1580-7.

71. Mkaouar, H., Mariaule, V., Rhimi, S., Hernandez, J., Kriaa, A., Jablaoui, A., Akermi, N., Maguin, E., Lesner, A., Korkmaz, B., and Rhimi, M. (2021). Gut Serpinome: Emerging Evidence in IBD. Int J Mol Sci 22. 10.3390/ijms22116088.

72. Li, S., Luo, L., Wang, S., Sun, Q., Zhang, Y., Huang, K., and Guan, X. (2023). Regulation of gut microbiota and alleviation of DSS-induced colitis by vitexin. Eur J Nutr 62, 3433–3445. 10.1007/s00394-023-03237-2.

73. Mikami, A., Ogita, T., Namai, F., Shigemori, S., Sato, T., and Shimosato, T. (2020). Oral Administration of Flavonifractor plautii, a Bacteria Increased With Green Tea Consumption, Promotes Recovery From Acute Colitis in Mice via Suppression of IL-17. Front Nutr 7, 610946. 10.3389/fnut.2020.610946.

