## Supplementary figures and images for "A microbiome meta-transcriptomics pipeline identifies a novel human neutrophil elastase inhibitor that protects the colonic epithelial barrier"

### Supplementary Figures

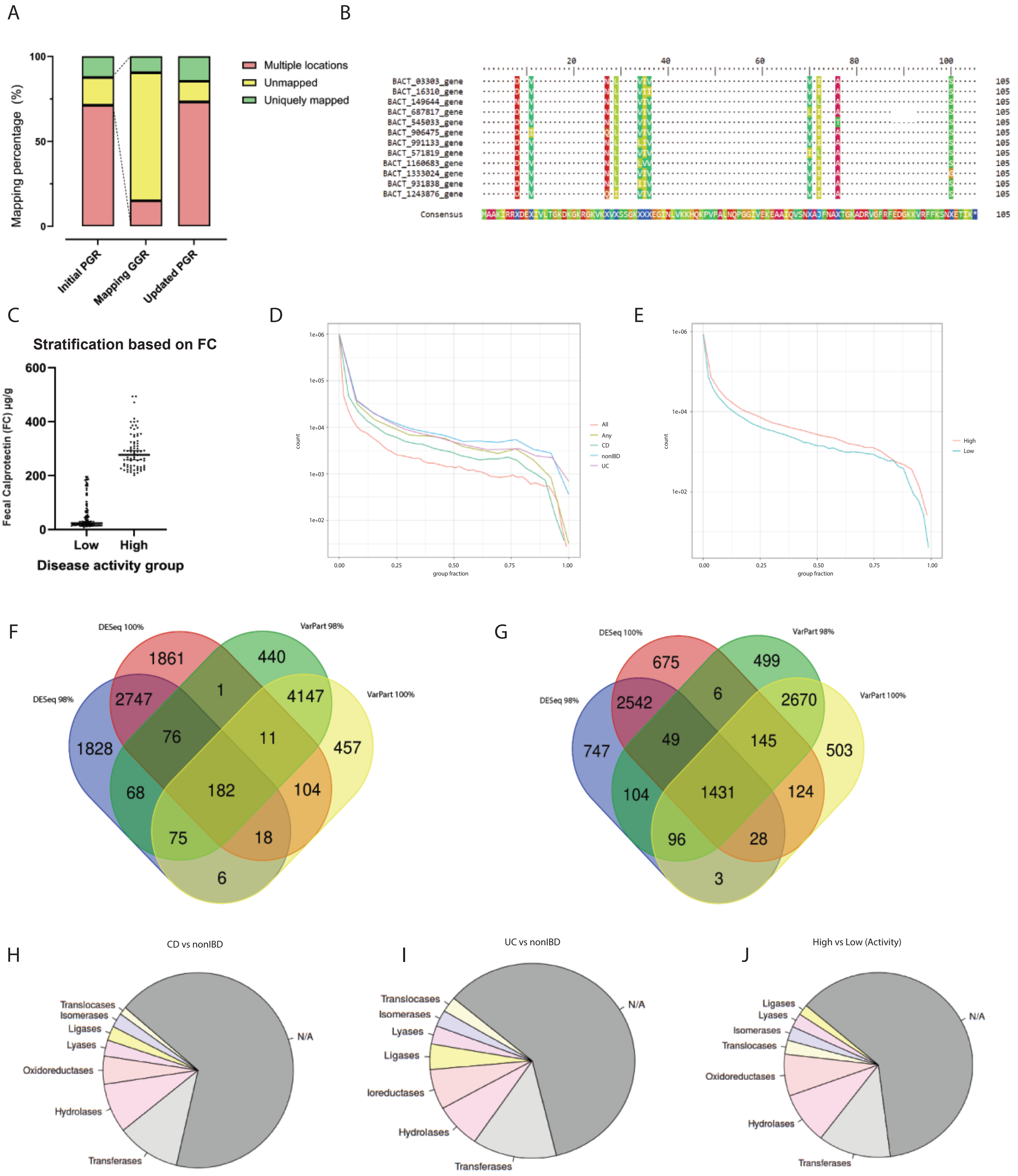

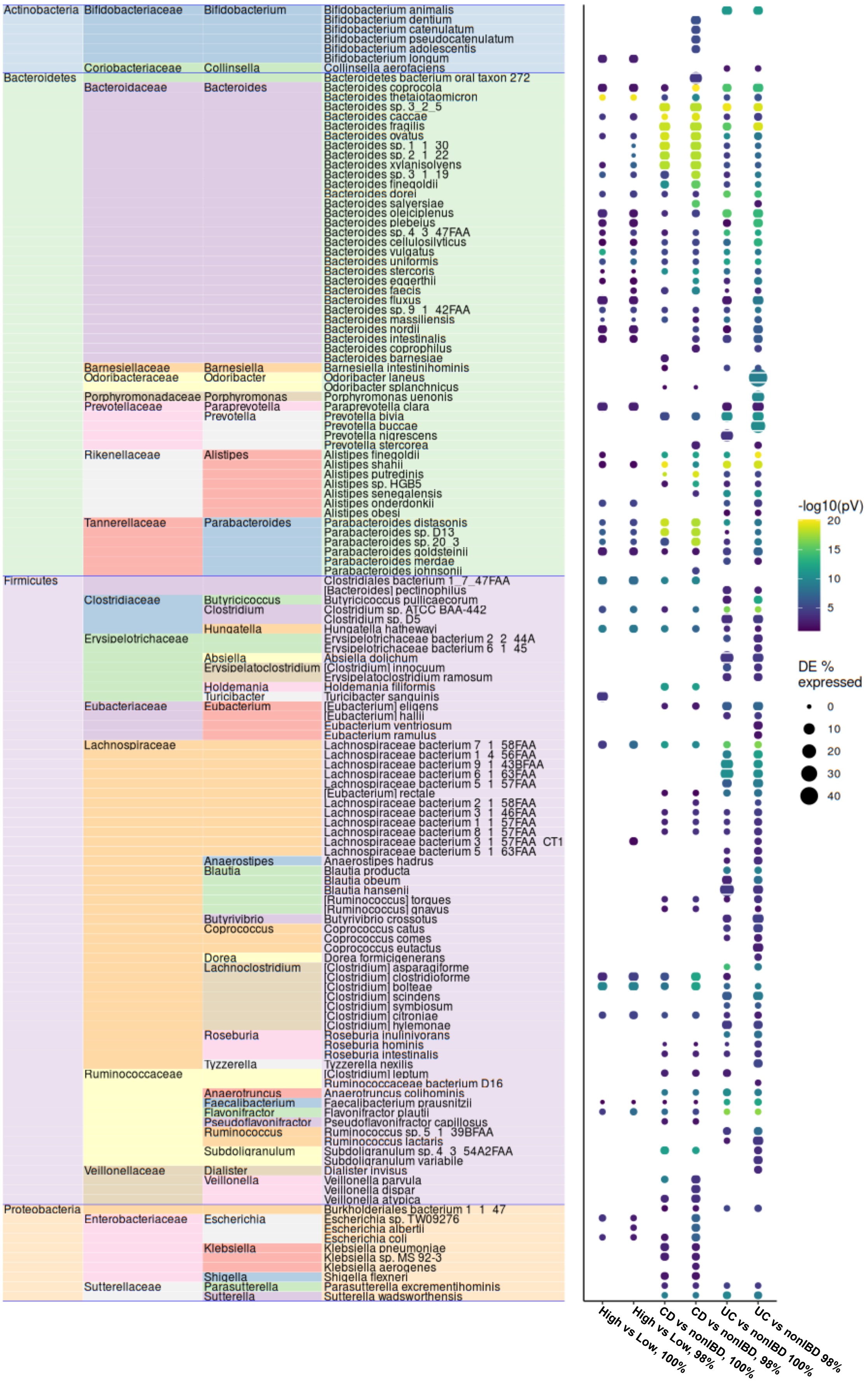

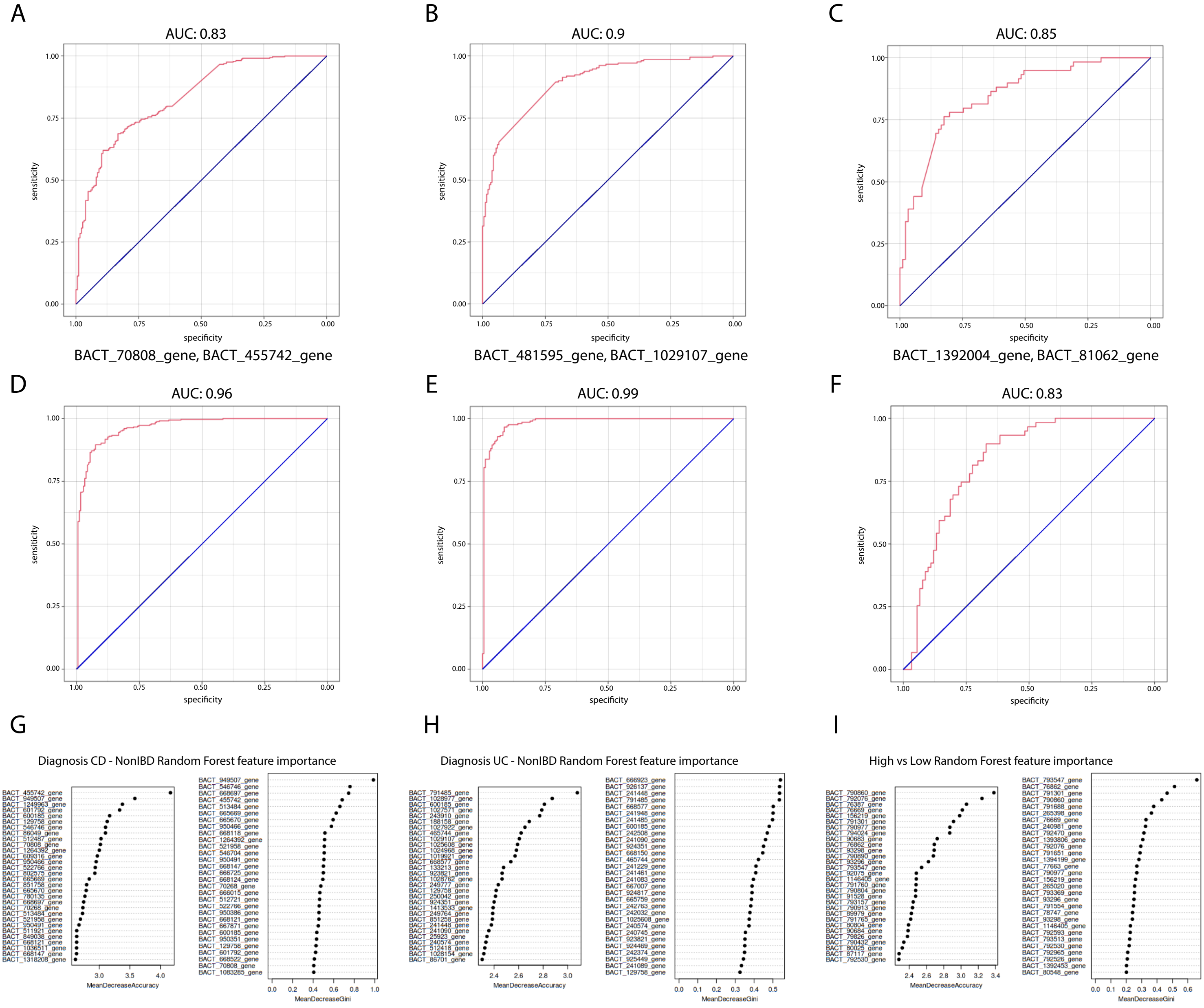

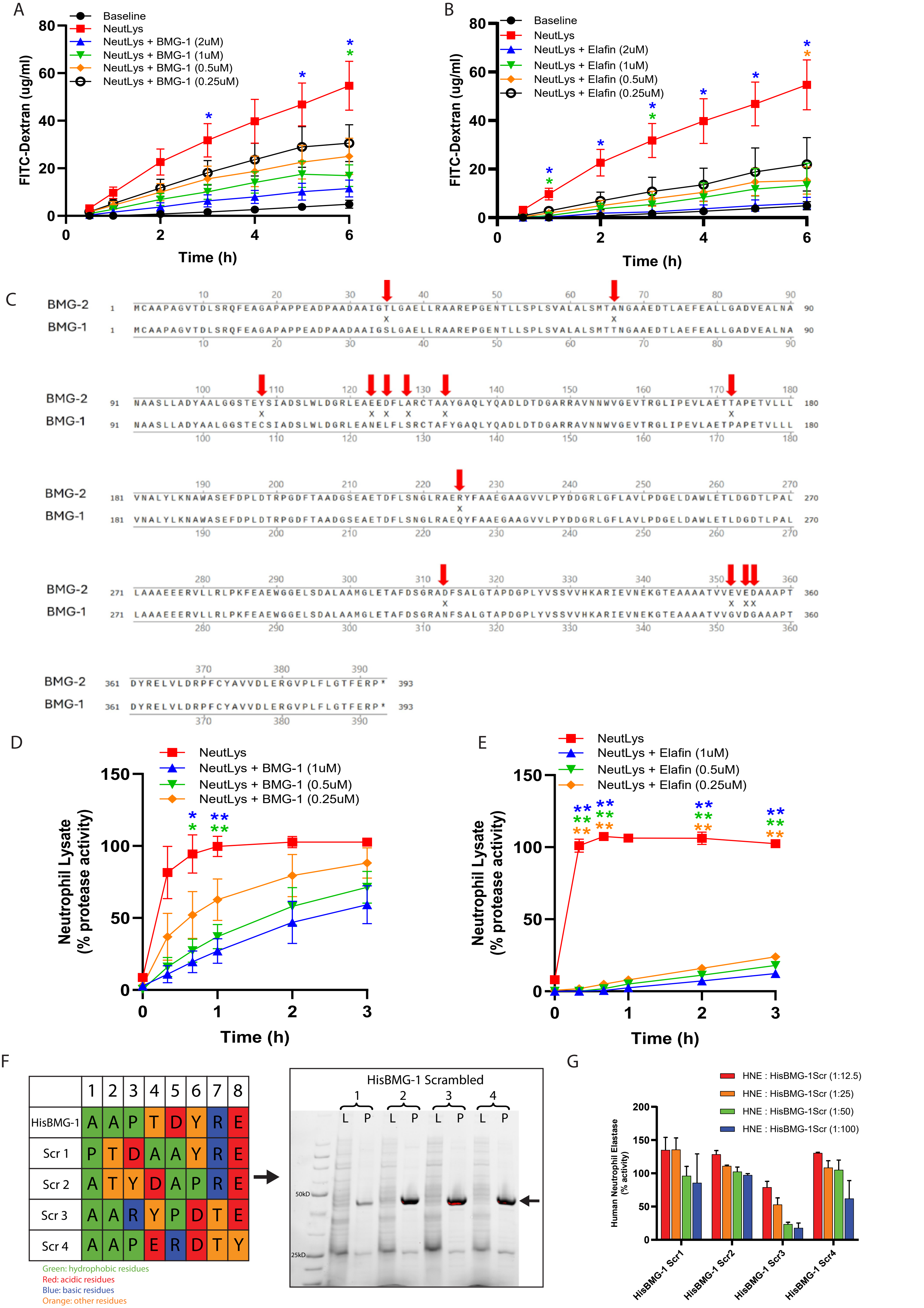

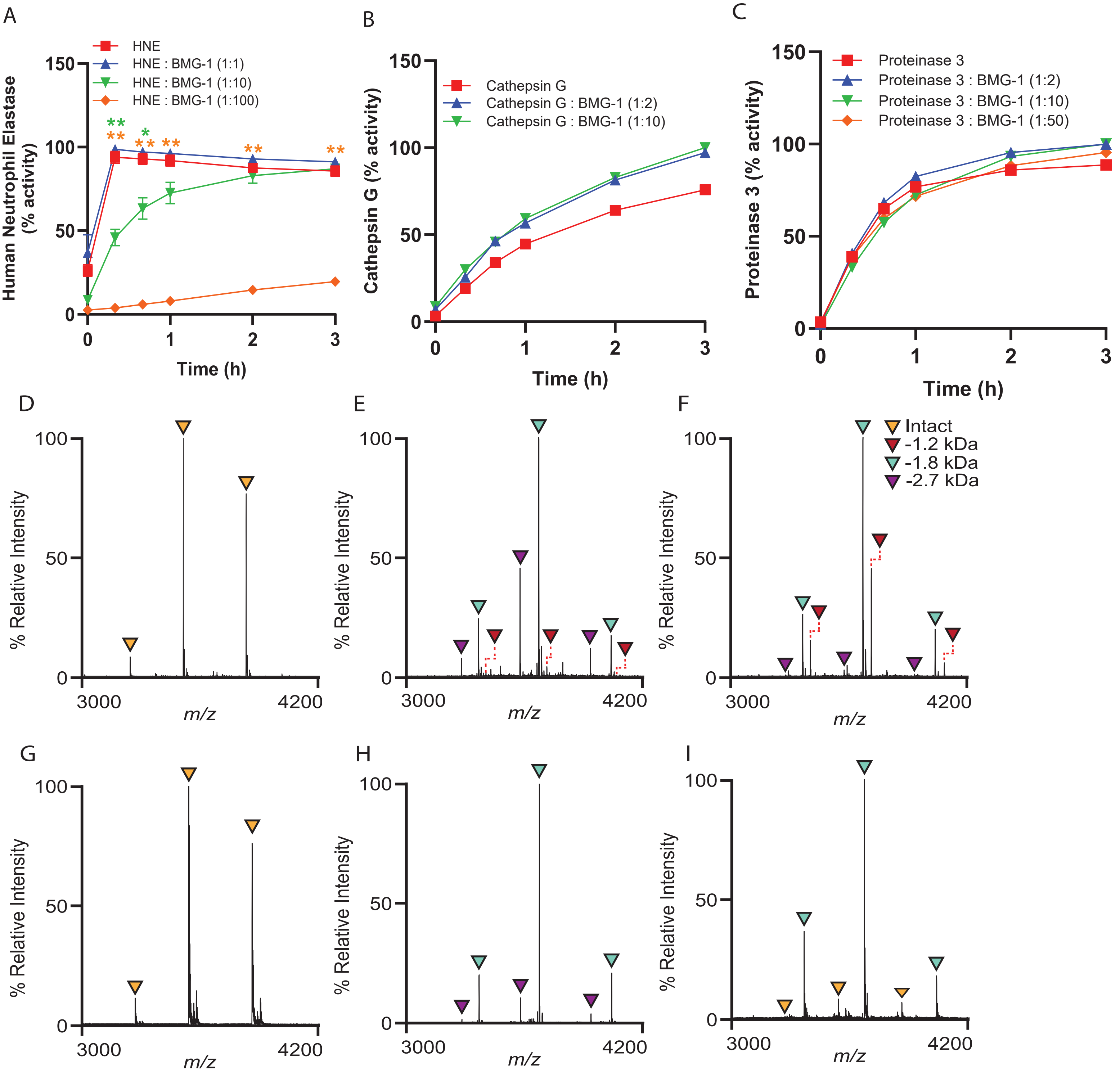

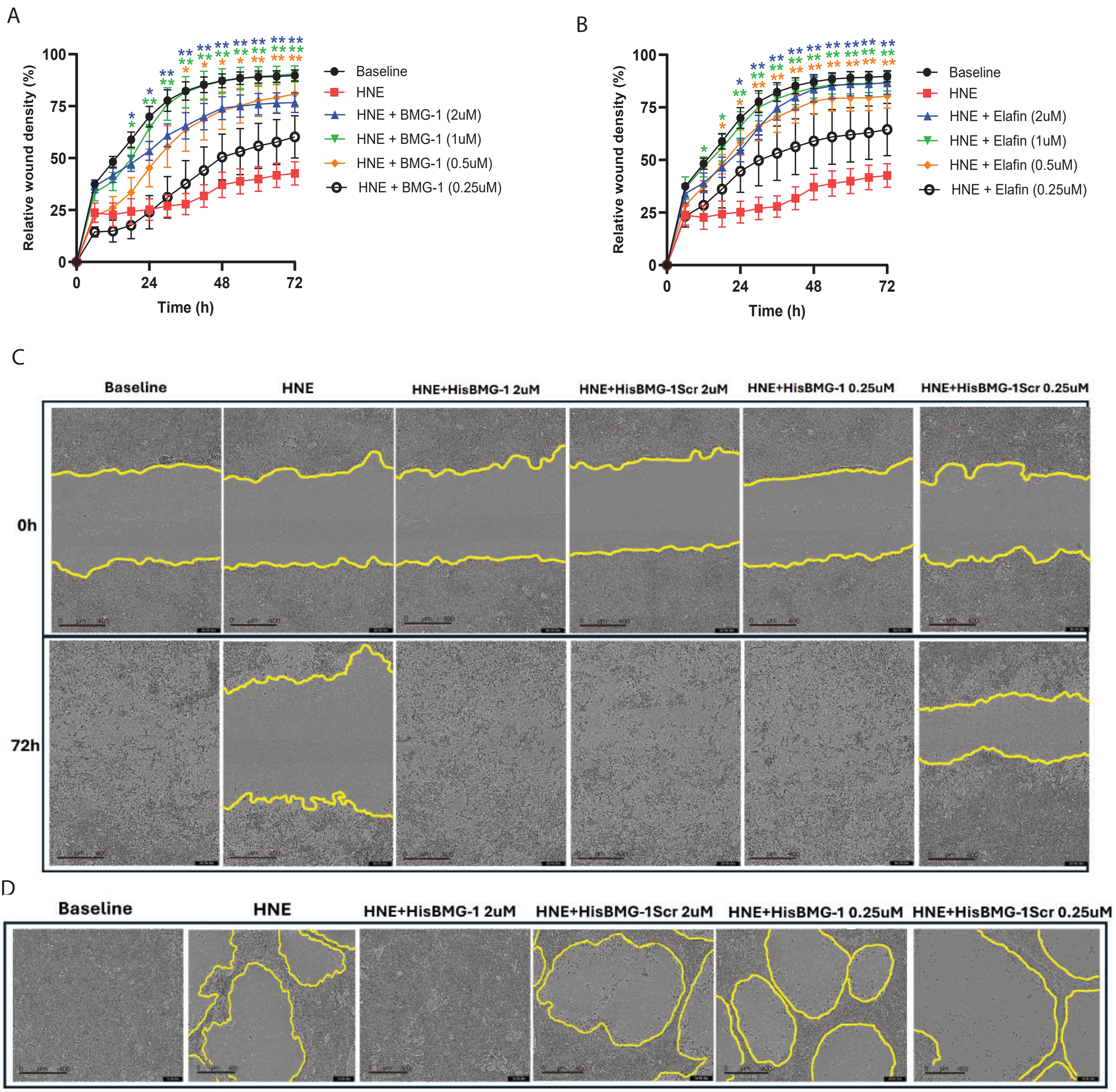
